# Vir2vec: A Genome-Wide Viral Embedding

**DOI:** 10.64898/2025.12.12.693901

**Authors:** Simone Rancati, Pablo Arozarena Donelli, Giovanna Nicora, Laura Bergomi, Tommaso Mario Buonocore, Micheal Aaron Sy, Sakshi Pandey, Mattia Prosperi, Marco Salemi, Riccardo Bellazzi, Christina Boucher, Enea Parimbelli, Simone Marini

**Author notes:** Equal contribution to first-authorship (Simone Rancati, Pablo Arozarena Donelli). Corresponding authors: Simone Marini; Enea Parimbelli.

## Abstract

Genomic language models (gLMs) enable the extraction of meaningful numerical representations from DNA sequences, yet existing approaches primarily focus on human genomes or limited viral datasets, and systematic evaluations of viral representations remain scarce. Here, we introduce Vir2vec, a family of decoder-only gLMs (17M–422M parameters) continually pretrained on a curated viral corpus comprising 565,747 complete genomes and genomic segment records assigned to 295 different species. We also present the viral Genome Understanding Evaluation (vGUE), a seven-task evaluation suite assessing viral representations across broad organism discrimination, evolutionary signatures, intra-genus species classification, variant subtyping, and host-associated phenotypes. Evaluated as a frozen feature extractor under nested cross-validation, Vir2vec achieves the highest balanced accuracy on three of seven tasks (full-genome virus detection, DNA-versus-RNA classification, and host-category prediction) and shares the highest performance on HIV-1 versus HIV-2 classification, while alternative models lead on the remaining three tasks. Taxonomic exclusion experiments further assess generalization of classification capabilities when an RNA-virus clade is excluded from pretraining at species and family levels. Finally, we introduce an argmax-routed SHAP explainability framework that maps feature attributions to codon-level sequence coordinates, highlighting receptor-binding-domain positions associated with SARS-CoV-2 human-versus-mouse host specificity discrimination. Together, Vir2vec and vGUE provide resources for advancing viral genomic representation learning and its systematic evaluation.

**Key Messages:**

- We present Vir2vec, a family of decoder-only viral genomic language models (17M, 138M, and 422M parameters) obtained via continual pretraining of Mistral-DNA on a curated, taxonomically restricted viral corpus (565,747 records, 295 species), generating fixed-length 4,096-dimensional embeddings for whole genomes and reads.
- We introduce vGUE, an initial evaluation suite for viral genomic foundation models spanning five biological dimensions: broad organism-level distinctions, genome-wide evolutionary fingerprints, intra-genus separation, lineage/subtype typing, and phenotypic context signal detection.
- Under a shared evaluation protocol (frozen embeddings plus shallow classifiers under nested cross-validation), Vir2vec leads on three of the seven vGUE tasks and ties for the best score on HIV-1 versus HIV-2, while Mistral-DNA or ModernBERT-virus perform better on the other three tasks.
- Taxonomic exclusion experiments demonstrate generalization of viral classification capabilities, with virus-versus-non-virus accuracies of 83.9% and 78.0%, and DNA-versus-RNA accuracies of 94.2% and 92.2%, following species- and family-level exclusions, respectively.
- We introduce an argmax-routed SHAP explainability framework that projects latent feature attributions back to single-codon resolution, highlighting receptor-binding-domain positions associated with human-versus-mouse sequence discrimination, including the region around codon 501.
- Context-truncated short reads (150-bp) and subtle intra-host tissue tropisms remain primary performance bottlenecks, while compact Vir2vec variants (17M and 138M) remain competitive on several benchmark tasks.

## Background

Viruses remain among the most pervasive threats to global health. For example, seasonal influenza causes around one billion infections and 3–5 million severe cases every year, leading to an estimated 290,000–650,000 respiratory deaths worldwide [1]; and dengue is endemic in more than 100 countries, with roughly half of the world’s population now living in at-risk areas, and an estimated 100–400 million infections annually [2]. Beyond headline pathogens, metagenomic surveys of environmental and human-associated microbiomes continue to uncover vast swath of viral diversity that has no close counterpart in current reference databases. Studies of marine and gut viromes, for instance, find that only a minority of viral reads can be confidently assigned to known taxa, highlighting how incomplete our picture of the global virosphere still is [3, 4, 5].

The COVID-19 pandemic dramatically accelerated genomic surveillance, turning high-throughput sequencing into a routine instrument of public health. However, this success has also exposed the limitations of the computational infrastructure sitting downstream of the sequencers. Traditional bioinformatics pipeline relies heavily on alignment to reference genomes or profile-based homology search, which are computationally intensive and intrinsically constrained by database coverage. Benchmarking efforts show that alignment-based methods and machine-learning methods perform well on near-reference sequences but degrade as genomes diverge or when working with short, noisy contigs in complex metagenomes [6, 7]. This is particularly problematic for viruses: reference collections are taxonomically biased, many clades remain sparsely sampled or entirely unrepresented, and viral taxonomy itself is in rapid flux. As a result, a substantial fraction of viral signal in clinical and environmental samples remains unclassified, limiting timely pathogen discovery and hampering quantitative ecological and epidemiological analyses [8, 9]. In parallel, advances in deep learning have begun to reshape computational genomics. Inspired by successes in natural language processing, several groups have proposed treating DNA as a sequence-based language and trained transformer-based foundation models that learn generic representations directly from nucleotide strings. A variety of architectures have emerged, ranging from decoder-only models capable of generating synthetic sequences [10, 11] to encoder-only models that produce numerical representations while preserving biologically meaningful signal. Among the encoder-based approaches, the Nucleotide Transformer (NT) family scales encoder-only models up to 2.5 billion parameters, pre-trained on human genomes supplemented with hundreds of additional species and achieves state-of-the-art performance across diverse genomics benchmarks [12]. DNABERT-2 similarly introduces a more efficient tokenizer and architecture than earlier genomic language models (gLMs), replacing fixed k-mer vocabularies with byte-pair encoding and enabling multi-species training at scale [13]. More recently, encoder–decoder architectures such as ENBED have demonstrated that long-context, byte-level transformers can be adapted to sequence-to-sequence tasks in genomics, further expanding the design space for genomic foundation models [14].

Despite these advances, existing gLMs are still only partially aligned with the needs of computational virology: the main architectures are not optimized for viral genomes, as training data are typically multi-species across kingdoms or primarily human-centric.

For example, Mistral-DNA-v1-422M-hg38 Genomic Language Model (gLM) is a 422M-parameter generative DNA model derived from the Mixtral-8×7B-v0.1 architecture [15], simplified for genomic applications by reducing the number of layers and the hidden size, and pretrained exclusively on 10 kb windows from the hg38 human genome assembly [16]. In addition, most published genomic foundation models are encoder-only architectures fine-tuned for specific classification or regulatory prediction tasks. As an example of such viral-focused efforts, the ModernBert-DNA-v1-37M-virus LLM is a 37M-parameter DNA model derived from ModernBERT [17], similarly simplified by reducing depth and hidden dimensionality, and pretrained on approximately 15,071 viral genomes longer than 1 kb, split into 1 kb segments [18]. While valuable, the diversity of pretraining datasets, genomic context lengths, and downstream evaluation protocols makes their relative strengths difficult to establish without matched comparisons. Recently, a handful of new models begun to develop the field of viral gLMs. ViraLM, for example, fine-tunes a DNABERT-2 backbone [13] on 50,000 viral genomes and shows improved virus-versus-non-virus classification in metagenomic contigs compared with classical approaches [19]. Other recent work has explored the use of large language models for hierarchical viral taxonomy prediction [20].

While promising, these approaches are typically optimized for a single downstream task or a narrow taxonomic slice and highlight the need to test whether reusable embeddings transfer across multiple viral groups and tasks. Therefore, a single, scalable embedding space that faithfully encodes many viral genomes across bacteria, archaea, plants, invertebrates and vertebrate hosts would provide a common numerical “language” on top of which heterogeneous tasks can be layered: virus vs non-virus detection in metagenomes; DNA vs RNA virus discrimination; host- or tissue-range prediction; lineage-level classification within high-priority pathogens such as SARS-CoV-2; and exploratory studies of sequence features associated with host range, subject to independent biological validation. To be practically useful, such an embedding engine must (i) be trained on a carefully curated, taxonomically broad, quality-controlled corpus; (ii) remain robust to unbalanced species distributions and heterogeneous genome lengths; and (iii) expose representations that are both geometrically well-structured (for clustering and visualization) and predictive in downstream supervised tasks.

To fill this research gap, here we introduce Vir2vec, a viral language model and embedding space developed to address the challenges detailed above. Vir2vec is obtained by continual pre-training of a 422-million-parameter, Mistral-style decoder-only transformer on a curated viral corpus reflecting the uneven taxonomic representation of publicly available genomic data, comprising more than 500,000 sequences associated with 295 reported species-level annotations. Using the tokenizer distributed with Mistral-DNA, Vir2vec learns context-rich nucleotide representations over this diverse corpus and exposes fixed-length, genome-level vectors via max-pooling over token embeddings.

Despite the growing interest in gLMs, existing benchmarking efforts offer limited coverage of viral representation tasks. Prior work spans a wide range of tasks and datasets, resulting in performance comparisons that are not always direct. With the Genome Understanding Evaluation (GUE), DNABERT-2 introduced the first effort to develop a standard benchmark for gLMs [13]. However, this benchmark covered only two virus-focused tasks, underscoring the lack of a standard evaluation framework for current and future viral gLMs. For this reason, we propose not only a new model, but also a task-based evaluation suite for viral representation learning: the viral Genomic Understanding Evaluation (vGUE) [13]. We systematically evaluate Vir2vec embeddings across a suite of downstream tasks that probe viral biology at multiple resolutions: (i) broad organism-level distinctions; (ii) genome-wide signatures and evolutionary fingerprint recognition; (iii) variant/subtype typing; (iv) intra-genus species distinction; and (v) phenotypic context signal identification.

Finally, we evaluated Vir2vec across two complementary applications: retention of broad classification signals under taxon exclusion, and localization of host-origin-associated attributions using SHAP [39] attributions paired with max-pooling argmax routing. These analyses examine retention of selected broad classification signals following taxon exclusion and identify sequence positions associated with host-origin classification, without directly assessing zoonotic risk or therapeutic targets.

By positioning Vir2vec as a viral-sequence embedding approach evaluated across multiple taxa—rather than a model tailored to a single dataset or endpoint—we aim to bridge the current gap between general genomic foundation models and narrow viral classifiers. Our results show that a single decoder-only transformer, trained on a curated viral corpus reflecting the uneven taxonomic representation of publicly available genomic data, can provide informative representations across several coarse-grained and lineage-level classification tasks. These findings highlight the potential of Vir2vec as a foundation for future computational virology applications, while emphasizing the importance of further validation across broader viral diversity.

## Methods

### Vir2vec architecture

Vir2vec builds upon the Mistral-DNA-v1-422M-hg38 model, a 422-million-parameter decoder-only transformer originally developed to model the human genome [16]. Each nucleotide sequence is first converted into discrete tokens using the tokenizer distributed with Mistral-DNA [16]. This tokenizer is based on byte-pair encoding (BPE) [21] adapted to DNA: rather than representing the sequence only as individual bases (A, C, G, T), it learns a vocabulary of short subwords that correspond to frequently occurring motifs and oligonucleotides in genomic data. This tokenization scheme allows the model to capture recurring sequence patterns in a compact way and to generalize more effectively across different genomes and host species [12, 22, 23].

Internally, input tokens are processed through model-specific hidden representations, which are subsequently projected into a 4,096-dimensional vocabulary-logit space. Sequence-level embeddings are obtained by applying max pooling across token positions. The base model incorporates modern efficient transformer components, including a mixture-of-experts (MoE) feed-forward design, grouped-query attention, and sliding-window attention (see Supplementary Note 1 for complete architectural details).

### Continual pretraining on viral genomes

To adapt the human-focused base model to the viral sequences represented in our corpus, we performed continual pretraining on a curated corpus of viral genomes [24, 25, 26, 27]. Starting from pre-trained Mistral-DNA weights, the model was further trained on hundreds of thousands of viral sequences across diverse taxonomic groups using a standard causal language modeling objective (predicting the next nucleotide-based token).

All layer dimensions, attention mechanisms, and network parameters were held identical to the original Mistral-DNA configuration during continual pretraining. Model selection was governed by tracking validation loss across epochs with early stopping to prevent overfitting. Following pretraining, the optimized Vir2vec network is used as a fixed feature extractor to generate 4096-dimensional embeddings for arbitrary viral sequences. Complete details regarding data preprocessing, hardware configuration, optimization schedules, and early-stopping criteria are provided in Supplementary Note 2.

### Data

Building a high-quality viral pretraining corpus requires systematic quality control and taxonomic balancing across heterogeneous public repositories. We compiled a curated viral corpus by integrating complete genomes from NCBI Virus [24] and BV-BRC [25], further enriched with specialized databases including GISAID [26], LANL-HIVdb [27], and HBVdb [28] (Figure 1A).

**Figure 1.**
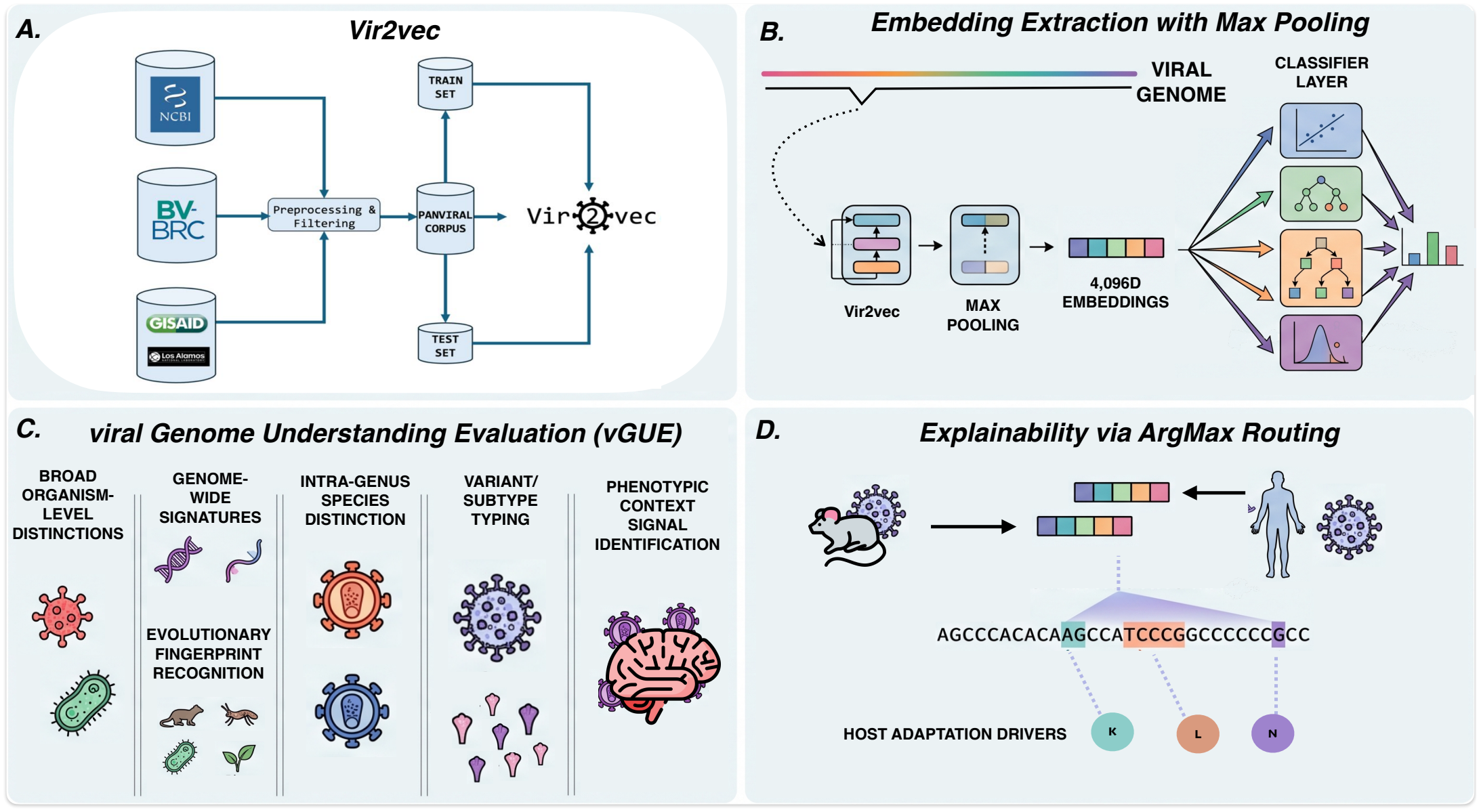
Vir2vec training workflow, embedding extraction mechanism, vGUE benchmark suite, and explainability framework.(A) High-quality complete viral genomes are compiled and filtered from public databases (NCBI, BV-BRC, and GISAID) into source-label-proportion-preserving train and test partitions to perform continual pretraining of the Vir2vec foundation model. (B) Nucleotide sequences (genomes, contigs, or short reads) are tokenized and processed through Vir2vec, followed by token-level max-pooling to extract fixed-length 4,096-dimensional embeddings for downstream classifier layers. (C) The viral Genome Understanding Evaluation (vGUE) benchmark assesses representation quality across five biological dimensions: broad organism-level distinctions, genome-wide evolutionary signatures, intra-genus species separation, variant/subtype typing, and phenotypic context signal detection. (D) Explainability via argmax routing transfers latent SHAP feature attributions directly back to nucleotide-based token coordinates to highlight sequence regions associated with SARS-CoV-2 human-versus-mouse host-origin classification. *Alt text: Four-panel schematic of data curation, Vir2vec embedding extraction, seven evaluation tasks grouped into five domains, and sequence-level attribution mapping*.

To ensure high sequence fidelity while preserving broad genomic diversity, candidate sequences were subjected to strict inclusion criteria regarding genome completeness and missing base thresholds. To mitigate the uneven taxonomic representation observed in publicly available viral genomic datasets, we implemented a two-step balancing strategy: filtering out under-represented or potentially misannotated entries while down-sampling highly represented taxa such as *SARS-CoV-2* and *Alphainfluenzavirus*.

The final curated corpus comprises 565,747 complete genome and genomic segment records distributed across 295 reported species annotations derived from the source databases, reflecting the diversity of viral taxa represented in the selected repositories. The dataset was split 70/30 into training and test partitions, preserving source taxonomic-label proportions across both subsets, with a small validation set drawn from the training split for hyperparameter tuning. The held-out test partition was kept strictly untouched during training and used exclusively to construct downstream evaluation benchmarks, preventing direct reuse of those records in pretraining, although sequence homology across partitions was not systematically quantified. Complete details regarding data collection and preprocessing are provided in Supplementary Note 3 and 4, while distributions of the final training corpus (taxons, hosts, and source repositories) are shown in Tables S1-S5.

### The vGUE Benchmark suite

The Genome Understanding Evaluation (GUE) benchmark, introduced alongside DNABERT-2 (Zhou et al., 2024) [13], marked the first attempt to standardize evaluation for genomic language models (gLMs). Compiling 36 datasets across nine task types, GUE primarily focuses on eukaryotic organisms (human, mouse, yeast). Its coverage of viral genomics, however, is restricted to two isolated tasks: SARS-CoV-2 variant classification and a broad species classification. Traditional eukaryotic benchmark tasks—such as promoter detection or epigenetic mark prediction—do not translate effectively to viral genomes, which feature high coding density, compact architectures, and a fundamental reliance on host machinery for replication. Consequently, broader genomic benchmarks do not explicitly test several viral sequence representation tasks of interest.

To address this gap, we introduce the viral Genome Understanding Evaluation (vGUE), a seven-task evaluation suite designed to probe gLMs on tasks adapted to viral biology (Figure 1C). Rather than serving as an alternative to optimized traditional bioinformatics algorithms (e.g., alignment tools like BLAST), vGUE is structured as a capability evaluation framework. Its goal is to compare frozen representations across tasks spanning broad taxonomic distinctions and sequence-associated phenotypes, providing an initial framework for systematic evaluation that can be further strengthened through standardised data splits and homology-aware validation protocols.

### Tasks

vGUE groups downstream evaluation tasks into five biologically motivated categories:

1. Broad organism-level distinctions: Distinguishing viral sequences from non-viral genomic backgrounds and identifying high-level taxonomic boundaries in both complete genomes and short-reads or fragmented sequencing data.
2. Genome-wide signatures and evolutionary fingerprints: Capturing global compositional biases and evolutionary constraints across whole genomes.
3. Intra-genus species distinction: Resolving fine-grained taxonomic relationships among closely related viral species.
4. Variant/subtype typing and phenotypic context signals: Identifying sub-species lineages and host-range determinants.
5. Phenotypic context signal identification: Capturing subtle adaptations of a same subtype virus to different environments within the host.

Instantiated across seven representative use cases (Table 1), vGUE spans full genomes and short reads, DNA and RNA viruses, diverse host taxa, and both taxonomic and phenotypic endpoints. This collection provides an initial comparative assessment, although near-ceiling performance on some tasks limits their discriminative power and further split and homology controls are needed. Description of all datasets used in the vGUE benchmark suite are included in Supplementary Note 5.

**Table 1.** Overview of the vGUE benchmark suite.

| Category | Task | Purpose | Data |
|---|---|---|---|
| Broad organism-level distinctions | Virus vs non-virus classification | Evaluate whether the model’s embeddings capture the fundamental features across full genomes that differentiate viral from cellular life. | Viral full genomes from corpus test set and 9,600 bacterial genomes from RefSeq fragmented in 5kb contigs. |
|  | Virus vs non-virus reads classification | Tests the model’s ability to recognize viral sequence signatures in a very limited context. | Real Illumina paired-end reads from NCBI BioProject PRJNA389927 [40]. 1,000 reads per class sampled from viral-enriched (SRR5665153) and microbial-enriched/non-viral (SRR5665119) runs. |
| Genome-wide signatures and evolutionary fingerprint recognition | DNA vs RNA classification | Probes whether the model has learned the deep evolutionary and genome-wide signatures that distinguish DNA-based viruses from RNA-based viruses. | 5,000 DNA and 5,000 RNA complete genomes from corpus test set. |
|  | Host Prediction | Evaluates whether the model captures host-associated genomic-wide signatures that allow assigning viruses to broad host categories. | 2,000 complete genomes from different families grouped in 4 classes based on the host: vertebrates, invertebrates, bacteria, and plants. |
| Intra-genus species distinction | HIV-1 vs HIV-2 classification | Evaluates the model’s ability to detect subtle genomic differences within a genus. | 7,248 HIV-1 and 94 HIV-2 complete genomes. |
| Variant/Subtype typing | SARS-CoV-2 Lineage Identification | Measures how well the model resolves very fine-grained sequence differences between closely related SARS-CoV-2 lineages that differ by only a small number of mutations across otherwise highly similar genomes. | 1,000 complete genomes across 7 different SARS-CoV-2 lineages (Alpha, Beta, Gamma, Delta, Lambda, Mu, Omicron). |
| Phenotypic context signal identification | HIV-1 Tropism Prediction | Tests if the model can capture subtle adaptation of a same subtype virus to different environments within the host. | HIV-1 subtype B env CDS: 764 brain seqs and 1,325 plasma seqs. |

Note: Tasks are grouped by biological level (broad organism-level distinctions, genome-wide signatures, intra-genus species discrimination, variant/subtype typing, and phenotypic context) and described in terms of their objective ("Purpose") and the corresponding dataset ("Data"), including full genomes and real sequencing reads drawn from our curated viral corpus and from public repositories. The specific tasks and datasets listed here are illustrative examples; vGUE is designed to be flexible, as long as evaluations cover the same high-level categories of viral genome understanding.

### Models

To systematically investigate how pretraining domain, model architecture, parameter scale, and context window affect viral representation learning, we evaluated a diverse suite of models on vGUE:

- Vir2vec (17M, 138M, 422M): To evaluate the effect of parameter scaling within the Vir2vec model family, we benchmark all three parameter sizes.
- Mistral-DNA (17M, 138M, 422M): Pretrained exclusively on human genomic DNA (hg38 assembly) [29], these models serve as direct out-of-domain baselines at parameter scales directly analogous to Vir2vec. Comparing Vir2vec against scale-matched Mistral-DNA models helps assess the contribution of viral continual pretraining, while downstream performance may still depend on classifier choice.
- ModernBERT-DNA-v1-37M-virus: Encoder-only architecture pretrained on 15,071 viral reference genomes segmented into 1kb chunks [18], providing an architecture- and segment-level baseline trained on viral data.
- Evo 1 (`Evo 1 8k base` and `Evo 1 131k base`): A state-of-the-art 7-billion parameter (7B) genomic foundation model trained on a massive corpus of 2.7 million prokaryotic and bacteriophage genomes (~300 billion nucleotides). [41] Because Evo 1 was trained with vastly greater computational resources and parameter scale than the rest of the models—and specifically excludes eukaryotic viruses—including both the 8k and 131k context length variants allows us to evaluate whether ultra-large model capacity, long-context processing with Hyena-based architecture [42] (rather than Transformer-based), and massive phage/prokaryotic pretraining can compensate for the absence of eukaryotic viral data when capturing viral genomic signals.

### Embedding Generation

Vir2vec and models included in the vGUE benchmark are evaluated as frozen feature extractors: given an input nucleotide sequence, the model produces a single fixed-length sequence embedding (Figure 1 B). Architecturally, each input sequence is tokenized, and the model outputs language model head vocabulary logits across positions. To derive a sequence-level summary, we apply max-pooling across the token dimension for each vocabulary feature without masking left-padded positions. We deliberately selected token-level max-pooling over simple mean-pooling because downstream tasks depend heavily on the presence of local, high-signal motifs (e.g., host- or lineage-associated genomic patterns), which become diluted when averaged across thousands of non-informative background positions. Subsequent averaging across sliding windows ensures that longer genomes do not artificially dominate downstream representations simply by contributing a higher number of windows. Experiments supporting pooling choices are described in Supplementary Note 6 and Figure S1.

For each benchmark task, input sequences (genomes, contigs, or reads) and associated labels are stored in tabular CSV files, preserving raw nucleotide strings without case normalization or sequence modification. To handle large sequence volumes without exhausting system memory, datasets are processed in a streaming fashion in chunks of *N* rows and dispatched in mini-batches for GPU inference. Tokenization is performed in left-padded mode with a maximum context length of 4,096 tokens; left-padding is applied to respect causal attention dynamics by ensuring real nucleotide tokens precede padding. Sequences exceeding 4,096 tokens are partitioned into overlapping sliding windows using a stride of 512 tokens.

For each window, we extract unmasked language model head logits and apply max-pooling across the token dimension to yield a *V*-dimensional window vector. To obtain a single embedding **e** for multi-window inputs, we compute the mean across window vectors:

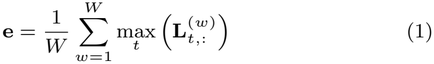

where *W* is the total number of windows, *t* indexes token positions within window *w*, and **L**^(*w*)^ represents the unmasked vocabulary logit matrix.

### Model-Specific Extraction Paradigms

While the pooling mechanism remains uniform across architectures, embedding dimensions and head mechanics reflect model-specific designs. For transformer-based causal decoders (Vir2vec and Mistral-DNA), output logits represent next-token distributions across a vocabulary space of *V* = 4,096. Conversely, the bidirectional encoder ModernBERT-DNA-v1-37M-virus produces 4,096-dimensional representations from its pretrained Masked Language Modeling (MLM) head, capturing non-causal motif predictions across all positions simultaneously. Finally, Hyena-based Evo models produce a 512-dimensional embedding without padding tokens, reflecting a native 512-character byte-level vocabulary rather than 4,096-token K-mer or BPE tokenizations. These representations are compared under a shared downstream probing protocol, while differences in architecture, prediction-head semantics and feature dimensionality may influence the observed rankings.

All forward passes are executed in full single-precision floating-point format (`fp32`) on a NVIDIA B200 GPU. Feature matrices and task metadata (label IDs, class names, and dataset metrics) are exported directly to uncompressed HDF5 archives for downstream evaluation.

### Classifier Layer

For all downstream tasks in vGUE, the pretrained sequence models are evaluated as frozen feature extractors. For each input sequence, we extract a single fixed-length sequence embedding **e** ∈ ℝ^*D*^ (where *D* = 4096 or *D* = 512 depending on the architecture). These representations serve as input features to shallow downstream classifiers trained independently per task and per model, enabling consistent comparisons of model representations under each classifier while highlighting the influence of representation–classifier combinations on predictive performance.

We select four modeling strategies spanning complementary inductive biases:

- Logistic Regression (LogReg): Operates with standard feature scaling (`StandardScaler`) and an *l*_2_-regularized L-BFGS solver to evaluate linear separability.
- Random Forest (RF): A non-parametric tree ensemble (*N*_trees_ = 300) capturing non-linear feature interactions.
- XGBoost (XGB): A histogram-based gradient-boosted decision tree classifier (`tree method=“hist”`) evaluating sequential gradient-boosted non-linear boundaries.
- Anomaly Detection Ensemble (AD): An unsupervised score-fusion ensemble combining a Nyström-kernelized One-Class SVM (OCSVM) and an Isolation Forest (IF) trained strictly on viral inliers. Base decision scores are *Z*-score standardized and linearly combined (*S*_ens_ = *w*_1_*Ŝ*_OCSVM_ + *w*_2_*Ŝ*_IF_), with mixing weights (*w*_1_, *w*_2_) and classification threshold *τ* jointly optimized out-of-fold across inlier quantiles to evaluate novel viral sequence boundaries.

Standard supervised classifiers (LogReg, RF, XGB) are applied across multi-class and phenotype prediction tasks, whereas Anomaly Detection is deployed specifically for virus versus non-virus discovery tasks.

Evaluation follows a nested stratified cross-validation setup. The outer loop performs 5-fold stratified cross-validation (80% training, 20% testing per fold) to estimate generalization performance. Within each outer training split, hyperparameter tuning (and decision threshold/weight calibration for AD) is conducted using an inner 3-fold stratified cross-validation via grid search, optimizing for macro-balanced accuracy. To handle class imbalance in supervised estimators, the hyperparameter grid explicitly evaluates both unweighted loss and sample-balanced class weighting (`class weight=‘balanced’`). The best hyperparameter configuration from the inner search refits the model on the outer training fold before evaluation on the held-out outer test fold.

Macro-balanced accuracy (BA) across outer test folds serves as the primary evaluation metric. For binary and multiclass tasks with *C* classes, BA is defined as the macro-average of per-class recall:

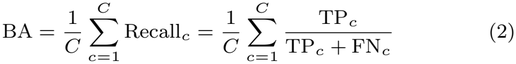

where TP_*c*_ and FN_*c*_ denote true positives and false negatives for class *c*, respectively. The supplementary tables present fold-level summaries of accuracy, balanced accuracy, precision, recall, Matthews Correlation Coefficient (MCC), and ROC-AUC.

Tasks derived from the curated corpus use the held-out partition; additional public-source tasks use separately retrieved sequences and accession-based exclusion where described in Supplementary Note 5, which does not by itself exclude closely related sequences. Precise descriptions of the datasets used in downstream tasks are in Supplementary Note 5.

### Generalization Analysis

A critical requirement for viral foundation models is the ability to generalize to novel, uncharacterized lineages. To examine retention of broad viral classification after excluding selected taxa from pretraining, we established a two-stage evaluation protocol using two pretraining data ablation regimes:

- Leave-Species-Out (ΔSpecies): Complete exclusion of all sequences belonging to the species *Enterovirus A* prior to pretraining.
- Leave-Family-Out (ΔFamily): Complete exclusion of the entire viral family *Picornaviridae* prior to pretraining, testing a broader taxonomic exclusion that includes the previously held-out species.

Embedding extraction for all sequences mirrored the primary vGUE pipeline by max-pooling unmasked *V* = 4,096 vocabulary logits across sliding windows, followed by window-level mean aggregation.

At first, primary downstream classifiers were trained using embeddings extracted from the held-out test sets of two core vGUE benchmark tasks, ensuring target clade sequences were strictly excluded: (i) binary Virus vs. Non-Virus (5 kb bacterial contigs) discrimination, and (ii) DNA vs. RNA classification. Both tasks were performed using histogram-based XGBoost classifiers (`xgb`) optimized via nested 5-fold outer and 3-fold inner stratified cross-validation. All fitted classifiers were subsequently frozen.

In a second phase, 1,000 genome sequences were randomly sampled from each excluded target taxon (*Enterovirus A* and *Picornaviridae*). Embeddings for these genomes were extracted using the corresponding ablated checkpoints and evaluated with the frozen downstream classifiers. For each exclusion setting, the evaluation comprised 1,000 withheld RNA-virus genomes and 1,000 negative control sequences, consisting of non-viral sequences for the virus-versus-non-virus task and DNA-virus genomes for the DNA-versus-RNA task. Classification performance was assessed using confusion matrices, accuracy, class-specific recall, and ROC-AUC computed from prediction scores.

### Host-Origin Explainability in SARS-CoV-2 S1

In viral genomics, predictive accuracy alone is insufficient if the underlying model remains a black box. Coupling genomic foundation models (gLMs) with model explainability translates latent representations into actionable biological insights, but attribution patterns alone cannot establish functional causality or rule out confounding and dataset-specific signals.

To examine sequence features associated with the host-origin labels, we constructed a binary classification dataset from the GISAID repository [26] comprising ~200 2, 055 bp genomic sequences of the SARS-CoV-2 *S*1 Spike gene, contrasting human-derived strains against mouse-adapted variants. Input sequences were tokenized and processed through *Vir2vec-422M* to extract token-level hidden states **H** ∈ ℝ^*N ×d*^. A regularized Logistic Regression classifier was then trained on sequence-level embeddings to predict host origin.

To preserve direct feature traceability back to sequence coordinates, we aggregate token representations using elementwise max pooling rather than standard mean pooling. While mean pooling averages representations and indiscriminately smears feature importance across all sequence tokens, max pooling preserves an exact spatial link. For each latent feature dimension *j*, max pooling retains the maximum activation alongside the specific token index that produced it, forming an *argmax routing map µ*(*j*):

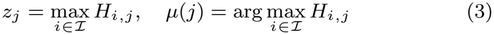

where *z*_*j*_ is the *j*-th feature of the sequence vector **z**, and *µ*(*j*) records the exact token origin of that dimension.

To measure feature importance, we calculate SHAP (SHapley Additive exPlanations) values *ϕ*_*j*_, which compute the marginal contribution of each latent feature *z*_*j*_ to the classifier’s prediction based on cooperative game theory. To our knowledge, this work introduces the first application of argmax-routed Shapley attributions to genomic language models. While existing gLM explainability frameworks overwhelmingly rely on attention-map heuristics, gradient-based saliency, or computationally expensive in-silico mutagenesis, our approach bridges latent game-theoretic feature attributions back to input sequence space. By leveraging the argmax routing map *µ*(*j*), each feature’s SHAP contribution *ϕ*_*j*_ is transferred directly to its winning token *t*_*i*_ without spatial dilution:

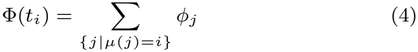

Finally, using tokenizer offset mappings, token attributions Φ(*t*_*i*_) are projected directly onto nucleotide sequence coordinates.

Rather than asserting ground-truth functional causation, this end-to-end pipeline explicitly reveals where the model focuses to distinguish host origins. This framework provides an interpretable methodology to generate hypotheses about host-associated sequence regions that require independent experimental and epidemiological validation.

## Results

### PCA evaluation of the embedding space

To obtain a global, task-agnostic assessment of how Vir2vec organizes viral diversity, we first analyzed its full 4,096-dimensional embeddings on the held-out viral corpus test set, comprising 30% of the corpus and stratified to preserve the proportions of the 295 retained source taxonomic labels. Using species labels only as a reference (i.e. treating each species as a putative cluster), we computed standard clustering quality metrics in the original embedding space. The silhouette [30] coefficient was slightly negative (−0.0115), whereas the Davies–Bouldin [31] and Calinski–Harabasz [32] indices were 3.0241 and 814.3, respectively. This combination of a near-zero silhouette with a relatively low Davies–Bouldin and high Calinski–Harabasz score does not support strong global species-level separation, although some local structure is visible in the low-dimensional projection. This is consistent with biological expectations: viruses frequently share conserved genes, replication strategies, and genome-wide constraints, so species are expected to lie along a continuum of similarity rather than forming perfectly isolated clusters (Figure 2A).

**Figure 2.**
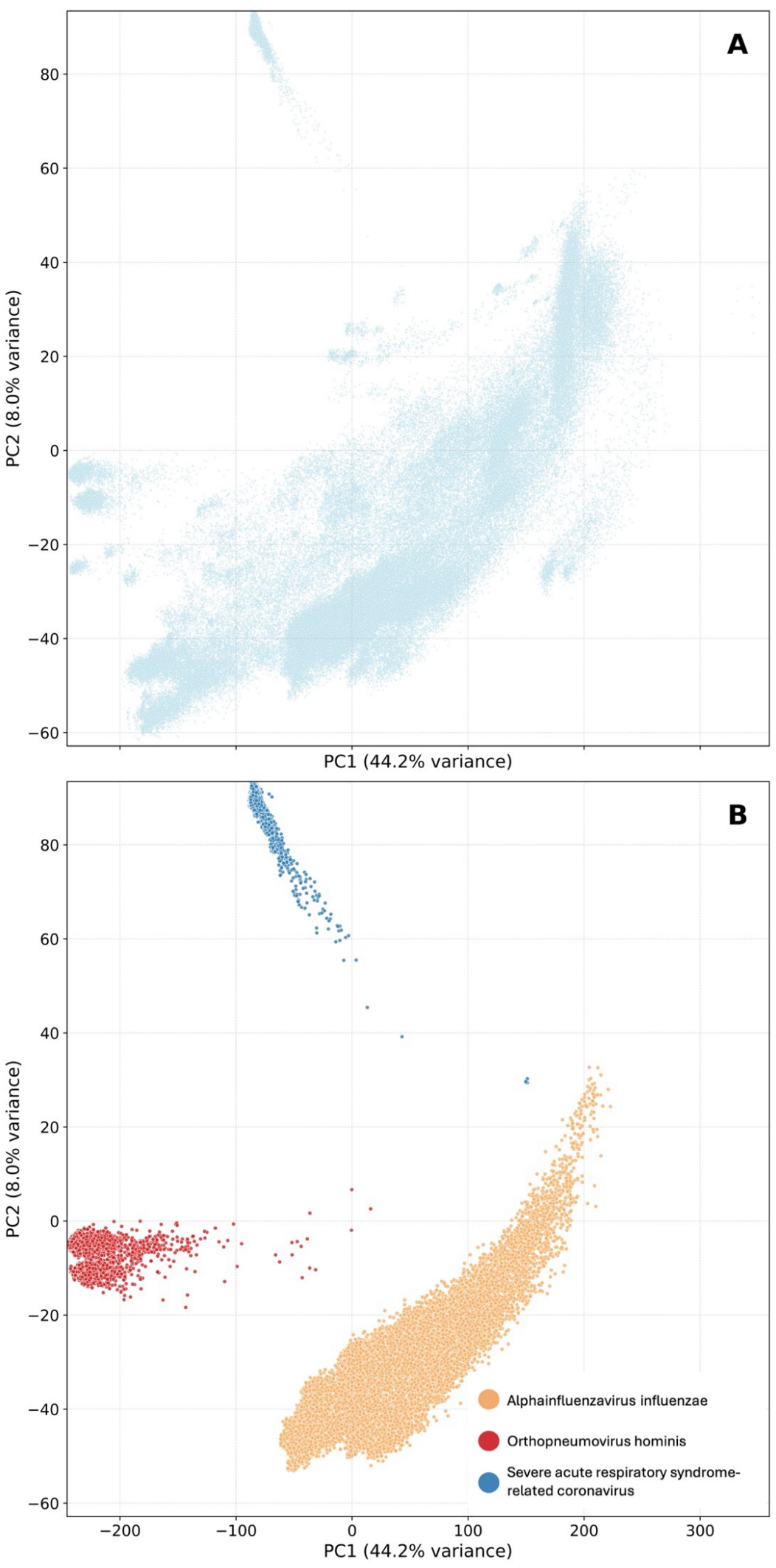
Global and species-specific geometry of the Vir2vec embedding space. (A) Two-dimensional PCA projection of the 4,096-dimensional Vir2vec embeddings for all 152,750 genomes in the held-out viral test set. Each point is a single genome, shown in a uniform color. The distribution exhibits partially compact substructures and a broad gradient along PC1, consistent with a continuum of viral diversity rather than perfectly isolated species clusters. (B) Same PCA space, highlighting three clinically relevant respiratory viruses—Alphainfluenzavirus influenzae, Orthopneumovirus hominis and Severe acute respiratory syndrome-related coronavirus—in distinct colors. In this illustrative low-dimensional projection, the three selected species show partial separation, indicating species-associated structure without establishing linear separability across the complete corpus. *Alt text: Two scatterplots of principal components: the full viral test set and three selected respiratory-virus groups highlighted in distinct colors*.

To obtain a more interpretable view, we examined the same PCA space restricted to three clinically important respiratory viruses: *Alphainfluenzavirus influenzae, Orthopneumovirus hominis* and *SARS-related coronavirus* (Figure 2B). Even in two dimensions, these three species appear partly separated in this illustrative two-dimensional projection. This qualitative pattern suggests that the frozen representation contains some species-associated information for these selected organisms, but does not establish species-level separation across the full corpus.

### Comprehensive Evaluation on the vGUE Benchmark

To evaluate the representation quality of Vir2vec, we benchmarked its three parameter scales (17M, 138M, and 422M) against a diverse suite of genomic language models: (i) Mistral-DNA-v1 (17M, 138M, 422M), pretrained on human genome sequences (hg38); (ii) Evo-1 (8k and 131k context lengths), a 7B general prokaryotic and phage foundation model; and (iii) ModernBERT-DNA-v1-37M-virus, an encoder pretrained on a limited set of segmented viral reference genomes. All models were evaluated by extracting frozen feature representations across seven downstream tasks in the vGUE benchmark paired with shallow classifiers. The summary of performances per task is illustrated in Figure 3, while Table 2 details the exact mean Balanced Accuracy (BA) and standard deviation across nested cross-validation folds in top performance classifiers.

**Table 2.**
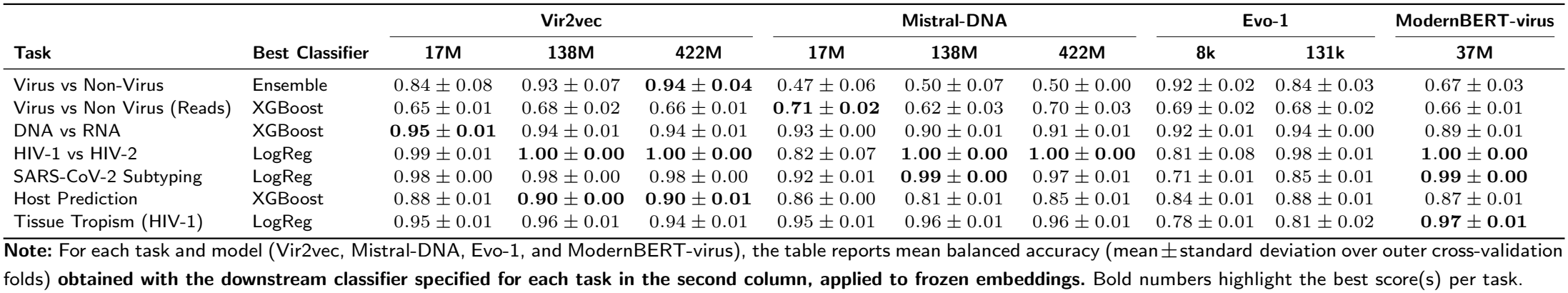
Performance comparison of genomic language models on the vGUE benchmark.

| Task | Best Classifier | Vir2vec |  |  | Mistral-DNA |  |  | Evo-1 |  | ModernBERT-virus |
|---|---|---|---|---|---|---|---|---|---|---|
|  |  | 17M | 138M | 422M | 17M | 138M | 422M | 8k | 131k | 37M |
| Virus vs Non-Virus | Ensemble | 0.84 ± 0.08 | 0.93 ± 0.07 | **0.94 ± 0.04** | 0.47 ± 0.06 | 0.50 ± 0.07 | 0.50 ± 0.00 | 0.92 ± 0.02 | 0.84 ± 0.03 | 0.67 ± 0.03 |
| Virus vs Non Virus (Reads) | XGBoost | 0.65 ± 0.01 | 0.68 ± 0.02 | 0.66 ± 0.01 | **0.71 ± 0.02** | 0.62 ± 0.03 | 0.70 ± 0.03 | 0.69 ± 0.02 | 0.68 ± 0.02 | 0.66 ± 0.01 |
| DNA vs RNA | XGBoost | **0.95 ± 0.01** | 0.94 ± 0.01 | 0.94 ± 0.01 | 0.93 ± 0.00 | 0.90 ± 0.01 | 0.91 ± 0.01 | 0.92 ± 0.01 | 0.94 ± 0.00 | 0.89 ± 0.01 |
| HIV-1 vs HIV-2 | LogReg | 0.99 ± 0.01 | **1.00 ± 0.00** | **1.00 ± 0.00** | 0.82 ± 0.07 | **1.00 ± 0.00** | **1.00 ± 0.00** | 0.81 ± 0.08 | 0.98 ± 0.01 | **1.00 ± 0.00** |
| SARS-CoV-2 Subtyping | LogReg | 0.98 ± 0.00 | 0.98 ± 0.00 | 0.98 ± 0.00 | 0.92 ± 0.01 | **0.99 ± 0.00** | 0.97 ± 0.01 | 0.71 ± 0.01 | 0.85 ± 0.01 | **0.99 ± 0.00** |
| Host Prediction | XGBoost | 0.88 ± 0.01 | **0.90 ± 0.00** | **0.90 ± 0.01** | 0.86 ± 0.00 | 0.81 ± 0.01 | 0.85 ± 0.01 | 0.84 ± 0.01 | 0.88 ± 0.01 | 0.87 ± 0.01 |
| Tissue Tropism (HIV-1) | LogReg | 0.95 ± 0.01 | 0.96 ± 0.01 | 0.94 ± 0.01 | 0.95 ± 0.01 | 0.96 ± 0.01 | 0.96 ± 0.01 | 0.78 ± 0.01 | 0.81 ± 0.02 | **0.97 ± 0.01** |

**Note:** For each task and model (Vir2vec, Mistral-DNA, Evo-1, and ModernBERT-virus), the table reports mean balanced accuracy (mean ± standard deviation over outer cross-validation folds) **obtained with the downstream classifier specified for each task in the second column, applied to frozen embeddings.** Bold numbers highlight the best score(s) per task.

**Figure 3.**
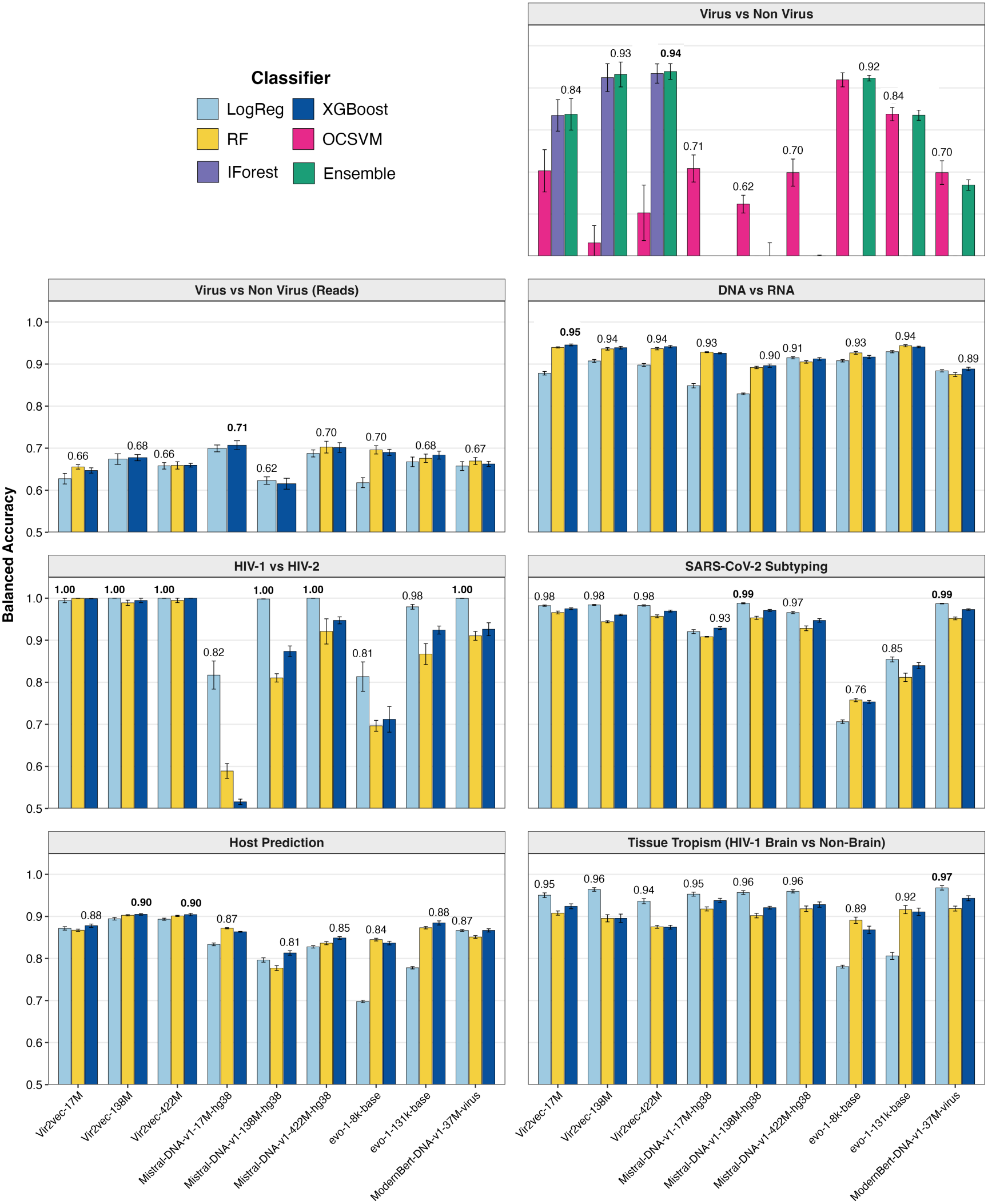
Performance of genomic language models across the viral Genome Understanding Evaluation (vGUE) benchmark. Evaluation of frozen sequence representations extracted from `Vir2vec` (17M, 138M, 422M), `Mistral-DNA` (17M, 138M, 422M), `Evo-1` (7B), and `ModernBERT-virus` (37M) across seven vGUE tasks. Downstream performance is measured using supervised shallow classifiers (LogReg, RF, XGBoost) for sequence-level discrimination, and unsupervised anomaly detection models (OCSVM, IForest, Anomaly Ensemble) for macro-level viral genome identification (*Virus vs Non-Virus*). Bars denote mean Balanced Accuracy across outer cross-validation folds, with error bars representing standard error (*±*SE). Numerical labels above the bars indicate the peak balanced accuracy achieved by each model per task. *Alt text: Seven grouped bar charts compare balanced accuracy by pretrained model and downstream classifier, with error bars*.

Broad Organism-Level Distinctions. For macro-level viral identification (*Virus vs Non-Virus*), models were evaluated under an anomaly detection framework using OCSVM, Isolation Forest, and Ensemble classifiers. Under this paradigm, Vir2vec achieves the highest result with the ensemble classifier, increasing from BA = 0.84 *±* 0.08 at 17M to a benchmark-leading **0.94** *±* **0.04** at 422M using the Anomaly Ensemble classifier. However, performance is classifier-sensitive: Mistral-DNA-17M reaches BA = 0.71 with OCSVM, compared with 0.70 for Vir2vec-17M and 0.60 for Vir2vec-422M under the same classifier. Thus, the ensemble results do not support a general claim that human-pretrained models fail to detect viral genomes. When evaluating short-read viral identification (*Virus vs Non-Virus (Reads)*), standard supervised classifiers (LogReg, RF, XGBoost) were applied instead of anomaly detection. Here, performance remains constrained across all architectures (BA = 0.62–0.71), suggesting that performance on the short-read dataset remains challenging across the evaluated representations, although the specific causes were not isolated.

Genome-Wide Signatures and Evolutionary Fingerprint Recognition. In genome type classification (*DNA vs RNA*), `Vir2vec-17M` achieves peak performance (0.95 *±* 0.01), outperforming larger human-trained and general foundation models. This suggests that compact Vir2vec representations capture signals associated with the DNA-versus-RNA virus labels in this dataset. For host range prediction (*Host Prediction*), `Vir2vec-138M` and `422M` achieve the highest overall accuracy (**0.90** *±* **0.00***/***0.01**) using XGBoost, consistently outperforming Mistral-DNA (0.81–0.86) and ModernBERT-virus (0.87), suggesting an advantage of the Vir2vec representations on the evaluated host-category task, without establishing a causal mechanism.

Intra-Genus Species Distinction, Variant Subtyping, and Phenotypic Context Identification. For fine-grained tasks—species separation (*HIV-1 vs HIV-2*), clade resolution (*SARS-CoV-2 Subtyping*), and tissue phenotype prediction (*Tissue Tropism*)—simple linear probes (LogReg) yield optimal performance, suggesting that the tested representations support effective linear discrimination on these particular datasets. While viral-pretrained models (Vir2vec and ModernBERT-virus) maintain near-ceiling accuracy across all model sizes (e.g., *≥* 0.98 on subtyping), naon-viral models exhibit a strong dependency on parameter scale: Mistral-DNA exhibits a marked improvement in performance between its 17M and 138M variants (0.82 *→* 1.00 on HIV species; 0.92 *→* 0.99 on subtyping). Conversely, Evo-1 achieves lower performance on the subtyping task (0.71–0.85), highlighting how differences in model architecture, pretraining data composition, context handling, and feature-extraction strategies may influence downstream classification performance.

Overall, these results show that viral continual pretraining can improve selected downstream evaluations, especially full-genome virus detection with the ensemble classifier, but the advantage is not consistent across all tasks and classifiers. Vir2vec attains the best reported balanced accuracy on three of seven tasks, shares the best result on HIV-1 versus HIV-2, and remains a useful point of comparison for viral sequence representation learning.

### Recognition under species- and family-level pretraining exclusions

We evaluated two classification tasks following exclusion of *Enterovirus A* at species level (ΔSpecies) and *Picornaviridae* at family level (ΔFamily) from pretraining. In each condition, 1,000 withheld RNA-virus genomes were evaluated against 1,000 negative-control sequences per task (Figure 4B).

**Figure 4.**
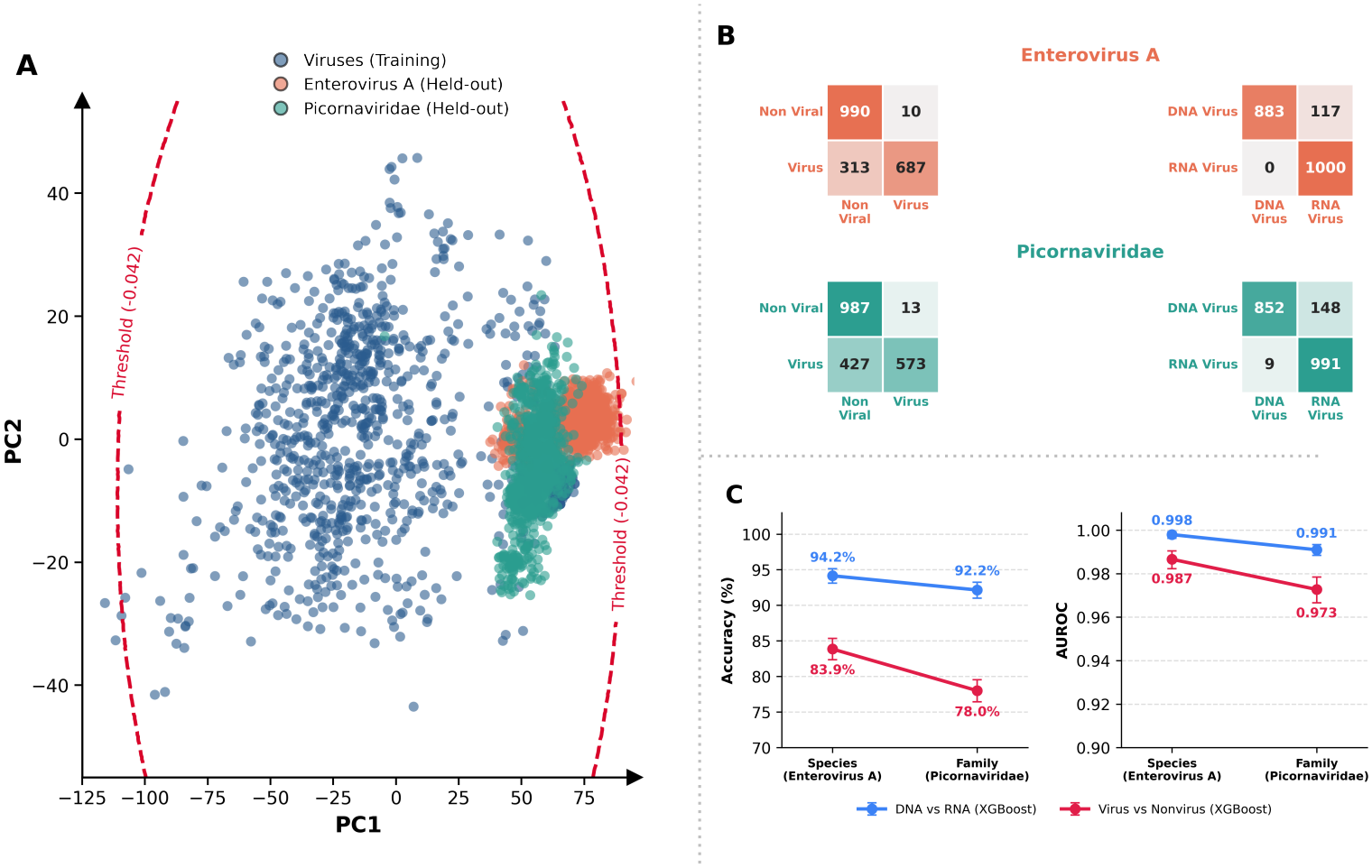
Classification after species- and family-level exclusion from Vir2vec-422M pretraining. (A) Two-dimensional PCA projection of baseline viral embeddings and target *Enterovirus A* (ΔSpecies) or *Picornaviridae* (ΔFamily) genomes. Dashed threshold guides are illustrative and should not be interpreted as the original high-dimensional decision surface. (B) Confusion matrices for virus-versus-non-virus and DNA-versus-RNA classification. Each exclusion setting includes 1,000 held-out target RNA-virus genomes and, respectively, 1,000 non-viral or 1,000 DNA-virus negative controls. Rows are true labels and columns are predicted labels. The virus-versus-non-virus confusion matrices are based on the anomaly detection ensemble, while the DNA-versus-RNA matrices are based on XGBoost. (C) XGBoost-based accuracy and AUROC in the resulting two-class comparisons: virus-versus-non-virus accuracy of 83.9% and 78.0% (AUROC 0.987 and 0.973), and DNA-versus-RNA accuracy of 94.2% and 92.2% (AUROC 0.998 and 0.991), for species- and family-level exclusions, respectively. These results concern the specified clade and comparator sets and do not establish generalization across multiple unseen families. *Alt text: PCA of withheld viral groups, four two-class confusion matrices, and plots comparing accuracy and AUROC under species and family exclusions*.

Virus-versus-non-virus classification. In the *Enterovirus A* exclusion condition, the confusion matrix contained 990 correctly classified non-viral controls, 10 false-positive controls, 687 correctly classified target viral genomes, and 313 false negatives. This corresponds to 83.9% accuracy and 68.7% viral sensitivity (99.0% specificity). For the *Picornaviridae* family exclusion, the respective counts were 987, 13, 573, and 427, corresponding to 78.0% accuracy and 57.3% viral sensitivity (98.7% specificity). Figure 4C reports corresponding AUROC values of 0.987 and 0.973.

DNA-versus-RNA classification. For the species-level exclusion, the classifier correctly recognised 883 of 1,000 DNA-virus controls and all 1,000 target RNA-virus genomes, corresponding to 94.2% accuracy, 88.3% DNA-virus recall, and 100.0% RNA-virus recall. Under family-level exclusion, it correctly recognised 852 DNA controls and 991 withheld RNA viruses, corresponding to 92.2% accuracy, 85.2% DNA-virus recall, and 99.1% RNA-virus recall. Reported AUROC values were 0.998 and 0.991, respectively (Figure 4C). As the class sizes are equal, accuracy and binary balanced accuracy coincide within rounding.

Embedding-space visualization. A PCA projection of baseline viral embeddings together with withheld RNA-virus targets is shown in Figure 4A. The projected grouping and dashed threshold guides are descriptive: a two-dimensional PCA plot does not reproduce the high-dimensional classifier decision surface or establish homology-independent recognition. Both exclusion settings target nested groups within a single RNA-virus family, so performance cannot be extrapolated to unseen DNA-virus families or to viral diversity more generally.

### Explainability Highlights Host-Origin-Associated Positions in SARS-CoV-2 S1

To examine sequence features associated with the host-origin labels, a regularized Logistic Regression classifier was evaluated on Vir2vec-422M embeddings across ~200 2,055 bp SARS-CoV-2 *S*1 Spike gene sequences using 5-fold nested cross-validation. The downstream classifier achieved robust host discrimination with a Balanced Accuracy of 98%. Following this classification analysis, token-level attributions were mapped to nucleotide sequence coordinates via our argmax-routed SHAP pipeline and aggregated into codon triplets to interpret the sequence features driving model focus.

Spatial Localization to the Receptor-Binding Domain. Analyzing signed SHAP profiles across the *S*1 gene reveals that predictive attributions are non-uniformly distributed across structural domains (Figure 5A), where negative SHAP values indicate feature contributions toward the mouse-adapted class. While the Signal Peptide (SP) and N-Terminal Domain (NTD) display relatively muted attribution scores, the Receptor-Binding Domain (RBD; ~ 1,000–1,650 bp, corresponding to aa 319–541) exhibits distinct, high-amplitude divergence between human and mouse cohorts. This concentration is consistent with the RBD carrying host-associated predictive information, but the attributions do not directly establish receptor-binding mechanisms or zoonotic risk [43, 44].

**Figure 5.**
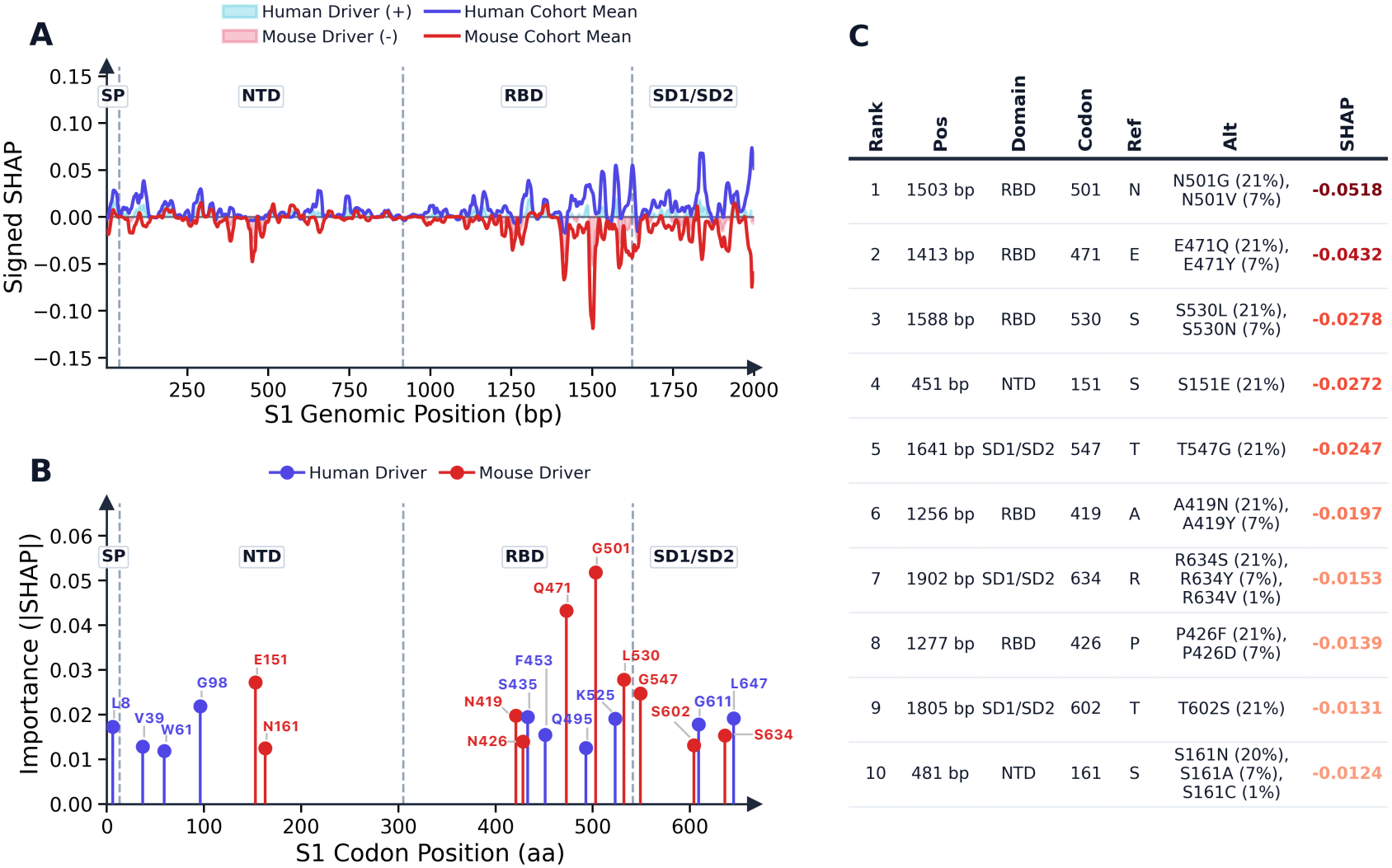
Explainability analysis highlights positions associated with SARS-CoV-2 human-versus-mouse host-origin classification in the *S*1 Spike gene. (A) Signed SHAP attribution profiles across *S*1 genomic coordinates (2,055 bp), comparing human-adapted cohort means (blue, positive SHAP) and mouse-adapted cohort means (red, negative SHAP) across Signal Peptide (SP), N-Terminal Domain (NTD), Receptor-Binding Domain (RBD), and Subdomains 1/2 (SD1/SD2). (B) Absolute SHAP importance (|SHAP|) per *S*1 codon position, highlighting top 10 human-associated residues (blue) and top 10 mouse-associated residues (red). (C) Ranked summary of the top 10 mouse-associated positions, detailing genomic coordinate (bp), domain, codon position, reference/alternate amino acid frequencies, and mean SHAP attribution scores. *Alt text: Profiles of signed SHAP attribution across the S1 gene, ranked residue contributions, and a table of the ten strongest mouse-associated sites*.

Identification of Known Murine ACE2 Adaptations. Aggregating feature attributions to amino acid positions reveals that the model concentrates on positions previously implicated in mouse adaptation (Figure 5B). Crucially, the most strongly attributed mouse-associated site lies at codon 501 (1,503 bp; SHAP = −0.0518), a residue position previously discussed in the literature on mouse adaptation (Figure 5C). This alignment matches established literature showing that mutations at position 501 in the Spike RBD enhance binding affinity to murine ACE2 (mACE2) during serial passage and natural host adaptation [43, 45, 46, 47].

Beyond codon 501, top predictive positions concentrate within RBD flexible loop regions (Figure 5B, C). The second-ranked site, codon 471 (1,413 bp; SHAP = −0.0432), corresponds to E471Q and E471Y substitutions within the receptor-binding motif. The third-ranked site, codon 530 (1,588 bp; SHAP = −0.0278), captures S530L and S530N mutations.

The correspondence between highly attributed positions and previously reported host-associated sites supports the biological plausibility of these candidate regions, but does not exclude dataset-specific signals or establish causal mechanisms.

## Discussion

Our findings demonstrate that genomic language models (gLMs) can provide useful numerical representations of viral sequences for selected downstream tasks. Across the vGUE benchmark, Vir2vec demonstrates that a single, frozen embedding space can capture complex biological signals ranging from broad viral-versus-non-viral discrimination to lineage typing and phenotype-associated signal detection. Continual pretraining on a curated viral corpus enables Vir2vec to learn transferable genomic representations across diverse viral classification tasks, providing a foundation for further evaluation across broader taxonomic diversity.

### Viral Continual Pretraining and Task-Dependent Strengths

A central observation from our benchmarking is that Vir2vec achieves leading results on selected tasks, particularly full-genome viral detection with the anomaly ensemble, DNA-versus-RNA classification, and broad host-category prediction; the advantage depends on task and downstream classifier. Pretraining on a curated corpus of 565,747 genome and complete-segment records across 295 source taxonomic labels provides a basis for learning sequence representations from the taxa covered by the training set, without establishing uniform coverage across viral diversity. This specialized pretraining yields top-tier performance across broad organism discrimination (BA = 0.94), DNA versus RNA classification (BA = 0.95), and multi-class host prediction (BA = 0.90). Crucially, Vir2vec achieves these results using simple, shallow classifiers operating on frozen 4, 096-dimensional representations, showing that useful predictive signals can be extracted without per-task fine-tuning of the language model.

Complementing the model itself, the vGUE evaluation framework provides a structured starting point for comparative evaluation in viral genomics. While existing benchmarks focus predominantly on eukaryotic regulatory tasks, vGUE systematically evaluates representation learning across five distinct levels of viral biology—spanning organism boundaries, evolutionary fingerprints, intra-genus species distinction, lineage subtyping, and phenotypic adaptation.

### Taxon-Exclusion Evaluation and Interpretability

A key question for viral sequence models is how prediction changes when taxonomic groups are omitted from pretraining. Our two nested ablations exclude *Enterovirus A* or the broader *Picornaviridae* family. Virus-versus-non-virus classification accuracy decreased from 83.9% to 78.0%, while DNA-versus-RNA classification accuracy decreased from 94.2% to 92.2% when moving from species-level to family-level exclusion. Similarly, viral-detection sensitivity decreased from 68.7% to 57.3%, whereas RNA-virus recall remained high at 100.0% and 99.1%, respectively. These findings support a restricted claim about performance on one withheld RNA-virus clade, in combination with the evaluated negative controls. They also provide a basis for investigating generalization across broader viral diversity through homology-aware evaluation protocols and more representative taxonomic sampling strategies.

Furthermore, coupling Vir2vec embeddings with max-pooling argmax routing and SHAP attributions provides a novel, transparent framework for sequence-level gLM interpretability. To our knowledge, this represents the first application of argmax-routed Shapley attributions to genomic language models, bypassing the spatial feature dilution inherent to standard pooling strategies. Applied to the SARS-CoV-2 *S*1 Spike gene, this pipeline mapped predictive signal directly back to genomic and amino acid coordinates, highlighting higher-magnitude attribution scores within the Receptor-Binding Domain (RBD; aa 319–541). Crucially, the highest-ranked mouse-associated attribution occurred near codon 501 (SHAP = −0.0518), alongside RBD loop mutations at positions 471 and 530. These results align directly with experimental literature establishing position N501 as a primary driver of high-affinity binding to murine ACE2 (mACE2), providing biological context for the observed attribution pattern without establishing a causal mechanism. Beyond retrospective validation, this interpretable gLM paradigm provides a scalable methodology for genomic epidemiology, offering a hypothesis-generating approach whose candidate sites would need experimental validation before any use in risk assessment or therapeutic design.

### Limitations and Bottlenecks

Despite these strengths, several methodological and biological limitations warrant consideration:

Contextualizing Short-Read Metagenomic Performance: Vir2vec was continuously pretrained on complete, full-length viral genomes to capture macro-genomic architecture and whole-sequence syntax. On short 150-bp sequencing reads (*BA* = 0.65–0.68), sequence-level frozen decoders naturally experience spatial context truncation. In contrast, specialized short-read classification tools in the literature such as DeepVirFinder [48] and RNN-VirSeeker [49], train dedicated 1D-CNNs or RNNs directly on short *k*-mer compositional frequencies. The higher balanced accuracy of `Mistral-DNA-17M` (0.71) relative to the Vir2vec variants (0.65–0.68) highlights a limitation of the current evaluation; the relative contributions of read length, sequence composition, and task-specific training remain untested.

Corpus Taxonomic Imbalance: Although hyper-dominant species (e.g., SARS-CoV-2 and Influenza) were down-sampled during corpus construction, the pretraining data remain strongly concentrated in a few taxa: the three most frequent species-level entries account for approximately 49% of the training sequences, based on the counts summarised in Tables S3 and S5. Only 295 retained source taxonomic labels are represented, including unresolved placeholders, with environmental and uncultured viral diversity underrepresented. Species-proportional sampling may transmit these imbalances to evaluation sets, inflating aggregate scores relative to taxon-macro-averaged performance.

Parameter Scale Efficiency: While scaling model capacity from 17M to 422M parameters yields modest marginal gains across frozen evaluation tasks (e.g., DNA vs. RNA classification peaking at 17M with 0.95*±*0.01; host range prediction stabilizing at 0.90*±*0.01), these results suggest diminishing gains with parameter scale on some of the tested tasks. The present experiments do not isolate the mechanism of this plateau or establish that compact models represent the full diversity of viral sequence space. The relationship between model scale, training composition, and feature-extraction strategy would require a controlled investigation. Compact variants (`Vir2vec-17M` and `Vir2vec-138M`) may offer useful performance–resource trade-offs for selected applications. Compared to multi-billion parameter foundation models such as Evo-1 (7B), compact Vir2vec variants achieve competitive or superior classification performance across several benchmark tasks while requiring substantially fewer model parameters than larger architectures.

Mutation-Level Signal Granularity Across High-Similarity Lineages: On downstream tasks involving near-identical sequences such as SARS-CoV-2 lineage subtyping (0.98–0.99) and HIV-1 versus HIV-2 species separation (0.99–1.00), Vir2vec exhibits performance parity with non-viral baselines (Mistral-DNA). These high scores may reflect limited task difficulty and potential sequence similarity within evaluation folds, rather than uniquely demonstrating transfer of genome-wide information. Additional homology-aware split analyses and less saturated tasks would be needed to distinguish genuine transfer from recognition of closely related sequences.

### Future Directions

Future developments for Vir2vec should focus on three primary avenues. First, the backbone can be specialized for metagenomic settings by mitigating taxonomic bias through continual learning on environmental viromes and uncultured contigs. To overcome performance limitations on short metagenomic reads, dedicated modules—such as multi-scale convolutional front-ends or assembly graph neural network aggregators—can be integrated to combine local motif context with global attention. Second, practical downstream applications can be expanded by deploying the argmax-routed SHAP explainability framework to investigate candidate sequence positions associated with host phenotypes, subject to functional validation, while utilizing the fixed-length 4,096-dimensional embeddings for exploratory genomic surveillance research after validation on independent datasets. Third, transitioning from frozen feature extraction to parameter-efficient fine-tuning (such as LoRA) will allow the transformer layers to capture subtle, localized mutational patterns critical for challenging phenotypic endpoints like tissue tropism and read classification.

## Conflicts of interest

The authors declare that they have no competing interests.

## Funding

This study has been funded in part by the NIH grant NIAID R01 AI170187; and by the Artificial Intelligence and Complex Computational Research Award, University of Florida.

## Data availability

The official source code for this study is available at https://github.com/pabloarozarena/Vir2vec. A versioned snapshot of the code has been archived in Zenodo (https://doi.org/10.5281/zenodo.24279702). The vGUE task datasets are available at https://huggingface.co/datasets/pabloarozarenad/vGUE-benchmark, and pretrained Vir2vec models are hosted at https://huggingface.co/pabloarozarenad/Vir2vec. Requests for controlled model access should include an institutional email address, a brief description of the intended use, and, where warranted by the proposed application, evidence of appropriate institutional oversight. Depending on the intended use, such oversight may include institutional biosafety, biosecurity, ethics, or human-subjects review.

## Acknowledgments

We gratefully acknowledge funding from the NIH and the University of Florida Artificial Intelligence and Complex Computational Research Award.

P. Arozarena Donelli is a PhD student enrolled in the National PhD program in Artificial Intelligence, XLI cycle, course on Health and life sciences, organized by Università Campus Bio-Medico di Roma.

Generative AI tools were used during the preparation of this manuscript for language refinement and manuscript editing. Specifically, ChatGPT (GPT-6, OpenAI) and Gemini (Gemini 4 Argon, Google) were used to improve linguistic clarity, readability, and overall presentation. The authors provided the instructions, critically reviewed all AI-generated suggestions, and retained full responsibility for the manuscript’s content, scientific accuracy, and final version.

## Authorship Contribution

All authors significantly contributed to the design of this study and the creation of the manuscript. SR, PAD, EP, SM, CB, RB and GN designed the study, SR and PAD wrote the first draft, that was revised by EP, GN, MS, RB, MP, CB, LB and TMB. SR, PAD,MAS, SP,LB and TMB performed the analysis under the supervision of RB, CB, MP, MS, EP, SM and GN. Funding acquisition was carried out by EP, SM and CB. All authors read and approved the manuscript in its current version.

## Notes

**Simone Marini, PhD,** is an Assistant Professor at the University of Florida, developing AI methods for computational genomics. **Enea Parimbelli, PhD,** is an Assistant Professor at the University of Pavia, specializing in biomedical informatics and AI-based clinical decision support.

### Competing Interest Statement

The authors have declared no competing interest.

### Summary of Updates

Increased the quality of the paper

