## Supplementary Figures and Tables for "Vir2vec: A Genome-Wide Viral Embedding"

*Briefings in Bioinformatics, 2026*, 2026, pp. 1–1

**Supplementary Note 1**

Vir2vec inherits its underlying neural architecture from the Mistral-DNA-v1-422M-hg38 base model [16], which adopts structural elements from the Mistral/Mixtral family of transformers [15] adapted for genomic sequences:

- **Tokenization:** Sequences are tokenized using a byte-pair encoding (BPE) vocabulary tailored for genomic data [21]. In addition to individual bases (A, C, G, T, N), the vocabulary encompasses frequent subwords and oligonucleotide motifs.
- **Feed-Forward Subnetworks:** In place of standard dense feed-forward blocks, each transformer layer contains a sparse Mixture-of-Experts (MoE) mechanism [15] that routes tokens to specialized expert subnetworks, scaling expressive capability without a linear increase in parameter computation.
- **Attention Mechanism:** Long-range context processing is facilitated via Grouped-Query Attention (GQA) and Sliding-Window Attention [15], reducing memory footprint during multi-kilobase context evaluation.
- Sequence-level Representation: For vGUE, max pooling is performed over the position-wise 4,096-dimensional vocabulary-logit outputs, giving a 4,096-dimensional feature vector; this output dimension should not be equated with the internal hidden-state width of every model variant.

**Supplementary Note 2**

Continual pretraining was conducted without modifying the underlying model architecture, keeping the total number of layers, attention heads, and hidden dimensions identical to the pre-trained Mistral-DNA base checkpoint.

- **Data Preprocessing:** Viral sequences were formatted into tabular datasets with raw nucleotide strings isolated under dedicated sequence fields. Inputs were tokenized and either padded or truncated to match the maximum context length supported by the base model. Training utilized standard unmasked causal language modeling.
- Optimizer & Learning Rate Schedule: Optimization was performed using AdamW [33] with an initial learning rate of $5 \times 10^{-4}$ and a weight decay coefficient of 0.01. A linear warmup schedule was applied over the first 1,000 gradient steps to ramp the learning rate from zero to $5 \times 10^{-4}$.
- **Batching & Precision:** Training was conducted in full 32-bit floating-point precision (fp32). The batch size was configured to 2 sequences per device with gradient accumulation over 4 steps, yielding an effective batch size of 8 sequences per parameter update. Gradient norms were clipped to a maximum threshold of 1.0 to ensure gradient stability.
- **Validation & Model Selection:** A dedicated validation split was evaluated at the conclusion of each epoch. Checkpoints were saved per epoch, and an early-stopping rule with a patience threshold of 3 consecutive non-improving epochs was enforced. The model checkpoint exhibiting the lowest validation loss across up to 10 total training epochs was selected as the final Vir2vec model.

**Supplementary Note 3**

Public viral repositories co-mingle high- and low-quality assemblies alongside variable metadata quality. To expose Vir2vec to natural, genome-wide structural constraints (such as gene order, codon usage, and long-range dependencies), sequence selection followed strict quality protocols:

- **Genome Completeness:** Only records representing complete viral genomes or complete genomic segments of segmented viruses were retained for pretraining, rather than fragmented reads or incomplete assemblies.
- Ambiguous Base Thresholds: Genomes were retained if they contained <1% ambiguous nucleotides (N) and no contiguous stretches exceeding 20 consecutive Ns. These limits ensure that at least 99% of bases per genome are fully resolved, reducing the proportion of unresolved input bases.
- **Taxonomic Harmonization:** Metadata fields across repositories were manually normalized to reconcile conflicting nomenclature (e.g., standardizing "HIV-1" vs. "Human immunodeficiency virus 1").
- **Deduplication**: Following harmonization, exact duplicate sequences **within and** across merged databases were removed.

**Supplementary Note 4**

Public repositories exhibit extreme sampling imbalances, simultaneously over-sampling prominent outbreak pathogens and including many incompletely resolved taxonomic labels.

- **Under-represented Taxa Filtering:** Taxa with fewer than 100 complete genomes were excluded, removing approximately 80,000 genomes. This reduced the number of retained source-database taxonomic labels from 28,229 to 295. These labels are not necessarily equivalent to ICTV-recognised species, and the filtering strategy favours well-sampled taxa.
- **Over-represented Taxa Down-sampling:** SARS-CoV-2 and *Alphainfluenzavirus* sequences were capped via random down-sampling to 100,000 genomes each. For influenza, down-sampling was stratified by genomic segment so that the segment distribution in the subsample directly mirrored the primary collection. Despite down-sampling, the three most frequent species-level annotations account for 192,731 of the 396,024 training records (48.7%; Tables S3 and S5).

The final collection of 565,747 viral genome and complete-segment records assigned to 295 retained taxonomic labels was structured into the following splits:

- **Training Split (70%):** Used for model pretraining. A minor validation subset was carved out from this split exclusively for hyperparameter selection and monitoring validation loss.
- **Test Split (30%):** Held out completely during pretraining and hyperparameter selection, used solely to construct datasets for downstream evaluation tasks. This separation prevents direct reuse of these records during pretraining, but does not by itself exclude highly similar sequences across partitions.

**Table S1. Taxonomic and metadata distribution of the Vir2vec training partition at the family level.** The 25 viral families with the highest number of sequences are reported, together with the median sequence length, most represented country and its proportion of sequences, collection-year range, proportion of sequences with an annotated genomic segment, and total number of sequences.

| Rank | Family | Median sequence length | Top Country | Collection years | Segmented sequences | Count |
|---|---|---|---|---|---|---|
| 1 | Orthomyxoviridae | 1743 | USA (60.4%) | 1900–2024 | 100.0% | 124774 |
| 2 | Coronaviridae | 29763 | USA (78.7%) | 1983–2024 | 0.0% | 72570 |
| 3 | Sedoreoviridae | 1351 | USA (13.3%) | 1961–2023 | 94.2% | 32363 |
| 4 | Retroviridae | 8958 | USA (23.8%) | 1984–2022 | 0.0% | 20508 |
| 5 | Flaviviridae | 10506 | USA (18.4%) | 1941–2024 | 2.2% | 19504 |
| 6 | Pneumoviridae | 15206 | USA (16.2%) | 1977–2024 | 0.0% | 13313 |
| 7 | Anelloviridae | 2740 | Tanzania (89.4%) | 1994–2023 | 0.0% | 11496 |
| 8 | Picornaviridae | 7361 | China (25.2%) | 1967–2024 | 0.0% | 9392 |
| 9 | Hepadnaviridae | 3215 | China (28.7%) | 1983–2024 | 0.0% | 9276 |
| 10 | Geminiviridae | 2739 | Brazil (12.0%) | 1990–2023 | 53.7% | 5748 |
| 11 | Circoviridae | 1768 | China (59.9%) | 2001–2024 | 0.0% | 4536 |
| 12 | Phenuiviridae | 3378 | China (75.6%) | 1969–2023 | 100.0% | 4003 |
| 13 | Spinareoviridae | 1996 | China (41.1%) | 1998–2023 | 100.0% | 3820 |
| 14 | Caliciviridae | 7509 | Japan (22.7%) | 1972–2024 | 0.0% | 3579 |
| 15 | Paramyxoviridae | 15347 | USA (29.0%) | 1969–2024 | 0.0% | 3281 |
| 16 | Togaviridae | 11482 | Brazil (38.6%) | 1905–2024 | 0.0% | 3076 |
| 17 | Rhabdoviridae | 11919 | USA (27.9%) | 1973–2024 | 0.0% | 2735 |
| 18 | Pospiviroidae | 359 | China (15.4%) | 1998–2024 | 0.0% | 2426 |
| 19 | Avsunviroidae | 337 | South Korea (6.8%) | 2005–2024 | 0.0% | 2307 |
| 20 | Poxviridae | 197185 | USA (31.1%) | 1902–2024 | 0.0% | 2260 |
| 21 | Nanoviridae | 1067 | India (32.5%) | 2008–2021 | 76.7% | 2036 |
| 22 | Potyviridae | 9786 | Japan (19.4%) | 1976–2023 | 0.0% | 1817 |
| 23 | Parvoviridae | 5010 | China (49.3%) | 2003–2024 | 0.0% | 1776 |
| 24 | Filoviridae | 18882 | Sierra Leone (53.2%) | 1976–2021 | 0.0% | 1700 |
| 25 | Papillomaviridae | 7905 | USA (15.2%) | 2000–2022 | 0.0% | 1634 |

**Table S2. Taxonomic and metadata distribution of the Vir2vec training partition at the genus level.** The 25 viral genera with the highest number of sequences are reported, together with the median sequence length, most represented country and its proportion of sequences, collection-year range, proportion of sequences with an annotated genomic segment, and total number of sequences.

| Rank | Genus | Median sequence length | Top Country | Collection years | Segmented sequences | Count |
|---|---|---|---|---|---|---|
| 1 | Betacoronavirus | 29766 | USA (80.1%) | 1987–2024 | 0.0% | 70949 |
| 2 | Alphainfluenzavirus | 1701 | USA (53.5%) | 1900–2024 | 100.0% | 70000 |
| 3 | Betainfluenzavirus | 1815 | USA (71.4%) | 1989–2024 | 100.0% | 52731 |
| 4 | Rotavirus | 1305 | Japan (16.2%) | 1975–2023 | 92.8% | 24599 |
| 5 | Lentivirus | 8960 | USA (24.6%) | 1987–2022 | 0.0% | 19825 |
| 6 | Orthoflavivirus | 10604 | USA (15.8%) | 1941–2024 | 0.0% | 15325 |
| 7 | Orthopneumovirus | 15209 | USA (16.0%) | 1977–2024 | 0.0% | 12796 |
| 8 | Orthohepadnavirus | 3215 | China (28.3%) | 1983–2024 | 0.0% | 9184 |
| 9 | Orbivirus | 1733 | South Africa (19.0%) | 1961–2023 | 98.8% | 7517 |
| 10 | Enterovirus | 7353 | China (27.1%) | 1970–2024 | 0.0% | 7355 |
| 11 | Circovirus | 1768 | China (61.1%) | 2001–2024 | 0.0% | 4420 |
| 12 | Betatorquevirus | 2799 | Tanzania (99.7%) | 2008–2016 | 0.0% | 4397 |
| 13 | Begomovirus | 2746 | Brazil (16.1%) | 2003–2023 | 72.0% | 4286 |
| 14 | Bandavirus | 3378 | China (86.9%) | 2000–2023 | 100.0% | 3301 |
| 15 | Hepacivirus | 9328 | USA (32.9%) | 1993–2022 | 0.0% | 3166 |
| 16 | Alphavirus | 11482 | Brazil (38.6%) | 1905–2024 | 0.0% | 3076 |
| 17 | Orthoreovirus | 2070 | China (24.0%) | 1998–2023 | 100.0% | 2824 |
| 18 | Norovirus | 7512 | Japan (26.1%) | 1972–2024 | 0.0% | 2460 |
| 19 | Pelamoviroid | 337 | South Korea (6.8%) | 2005–2024 | 0.0% | 2307 |
| 20 | Lyssavirus | 11923 | USA (22.1%) | 1973–2024 | 0.0% | 2230 |
| 21 | Orthopoxvirus | 197189 | USA (33.4%) | 1902–2024 | 0.0% | 2099 |
| 22 | Potyvirus | 9786 | Japan (19.4%) | 1976–2023 | 0.0% | 1817 |
| 23 | Babuvirus | 1075 | India (38.2%) | 2008–2021 | 72.6% | 1735 |
| 24 | Orthoebolavirus | 18882 | Sierra Leone (53.2%) | 1976–2021 | 0.0% | 1700 |
| 25 | Pospiviroid | 359 | South Korea (5.3%) | 1998–2023 | 0.0% | 1479 |

**Table S3. Source species-field annotation distribution in the Vir2vec training partition.** The 25 most frequent source species labels are shown with median length, main country, collection-year range, segmented-sequence proportion and total count; some annotations are unresolved placeholders at higher taxonomic levels and must not be interpreted as verified species.

| Rank | Source species-field label | Median sequence length | Top Country | Collection years | Segmented sequences | Count |
|---|---|---|---|---|---|---|
| 1 | Alphainfluenzavirus influenzae | 1701 | USA (53.5%) | 1900–2024 | 100.0% | 70000 |
| 2 | Severe acute respiratory syndrome-related coronavirus | 29764 | USA (81.0%) | 2008–2024 | 0.0% | 70000 |
| 3 | Betainfluenzavirus influenzae | 1815 | USA (71.4%) | 1989–2024 | 100.0% | 52731 |
| 4 | Rotavirus A | 1317 | Japan (16.0%) | 1975–2023 | 100.0% | 22568 |
| 5 | Human immunodeficiency virus 1 | 8950 | USA (21.4%) | 1994–2022 | 0.0% | 18698 |
| 6 | Caudoviricetes sp. * | 29165 | USA (99.4%) | 2010–2020 | 0.0% | 15578 |
| 7 | Orthopneumovirus hominis | 15209 | USA (16.0%) | 1977–2024 | 0.0% | 12796 |
| 8 | Orthoflavivirus denguei | 10539 | Thailand (16.6%) | 1956–2024 | 0.0% | 10440 |
| 9 | Hepatitis B virus | 3215 | China (28.3%) | 1983–2024 | 0.0% | 9184 |
| 10 | Bluetongue virus | 1659 | France (20.6%) | 1972–2023 | 100.0% | 5109 |
| 11 | TTV-like mini virus | 2799 | Tanzania (99.7%) | 2008–2016 | 0.0% | 4397 |
| 12 | Torque teno midi virus | 2597 | Tanzania (99.7%) | 2017–2017 | 0.0% | 3685 |
| 13 | Bandavirus dabieense | 3378 | China (86.9%) | 2000–2023 | 100.0% | 3301 |
| 14 | Hepacivirus hominis | 9328 | USA (33.6%) | 1993–2022 | 0.0% | 3093 |
| 15 | Enterovirus A | 7395 | China (49.7%) | 1998–2024 | 0.0% | 2736 |
| 16 | Norwalk virus | 7512 | Japan (26.1%) | 1972–2024 | 0.0% | 2460 |
| 17 | Circovirus porcine2 | 1767 | China (74.2%) | 2002–2024 | 0.0% | 2341 |
| 18 | Torque teno virus | 3140 | Tanzania (95.8%) | 1994–2021 | 0.0% | 2314 |
| 19 | Orthoflavivirus nilense | 10787 | USA (70.8%) | 1988–2023 | 0.0% | 2246 |
| 20 | Chikungunya virus | 11309 | Brazil (53.1%) | 1953–2024 | 0.0% | 2230 |
| 21 | Peach latent mosaic viroid | 337 | South Korea (6.8%) | 2005–2024 | 0.0% | 2220 |
| 22 | Bacteriophage sp. * | 65806 | Japan (88.2%) | 2010–2020 | 0.0% | 2195 |
| 23 | Lyssavirus rabies | 11923 | USA (23.4%) | 1973–2024 | 0.0% | 2106 |
| 24 | Monkeypox virus | 197190 | USA (34.4%) | 2001–2024 | 0.0% | 2012 |
| 25 | Orthoebolavirus zairense | 18882 | Sierra Leone (53.2%) | 1976–2021 | 0.0% | 1700 |

Table S3 note. * "Caudoviricetes sp." and "Bacteriophage sp." are unresolved source metadata placeholders, not formal species names. They are displayed for auditability of the original species field, not counted as independently validated species. Accordingly, the count of 295 refers to retained source taxonomic labels, not confirmed ICTV species.

**Table S4. Host distribution in the Vir2vec training partition.** The 25 most frequently represented host entries are shown with their corresponding NCBI Taxonomy ID and sequence counts. The table highlights inconsistencies in host annotation across database records, including multiple entries referring to the same host (e.g., Homo sapiens and human-related labels) and missing Taxonomy IDs for some host annotations.

| Rank | host_organism | host_taxid | Count |
|---|---|---|---|
| 1 | Homo sapiens | 9606.0 | 186638 |
| 2 | Homo sapiens | NA | 55006 |
| 3 | NA | NA | 33678 |
| 4 | human metagenome | 646099.0 | 13019 |
| 5 | Sus scrofa | 9823.0 | 12875 |
| 6 | Gallus gallus | 9031.0 | 4927 |
| 7 | Bos taurus | 9913.0 | 3989 |
| 8 | swine | NA | 3856 |
| 9 | chicken | NA | 2812 |
| 10 | Prunus persica | 3760.0 | 2249 |
| 11 | Human | NA | 2233 |
| 12 | Solanum lycopersicum | 4081.0 | 1895 |
| 13 | Anas platyrhynchos | 8839.0 | 1875 |
| 14 | Anatidae | 8830.0 | 1758 |
| 15 | Vitis vinifera | 29760.0 | 1469 |
| 16 | Canis lupus familiaris | 9615.0 | 1432 |
| 17 | duck | NA | 1289 |
| 18 | Equus caballus | 9796.0 | 1086 |
| 19 | Nicotiana benthamiana | 4100.0 | 1073 |
| 20 | Ovis aries | 9940.0 | 997 |
| 21 | Salmo salar | 8030.0 | 939 |
| 22 | Sus scrofa domesticus | 9825.0 | 918 |
| 23 | Solanum tuberosum | 4113.0 | 865 |
| 24 | Macaca mulatta | 9544.0 | 856 |
| 25 | Capsicum annuum | 4072.0 | 839 |

**Table S5. Sequence distribution across source viral repositories in the training partition of Vir2vec.** The percentage under the "Number of sequences" column indicates the proportion of sequences in the training partition (396,024 records), whereas the percentages listed for the "Top 3 species" represent the proportion of each species relative to all sequences originating from that specific repository.

| Source | Number of sequences | Top 3 species |
|---|---|---|
| NCBI Virus | 271,222 (68.49%) | Severe acute respiratory syndrome-related coronavirus (25.81%) |
| | | Alphainfluenzavirus influenzae (12.23%) |
| | | Betainfluenzavirus influenzae (7.85%) |
| BV-BRC | 98,462 (24.86%) | Alphainfluenzavirus influenzae 37.41%) |
| | | Betainfluenzavirus influenzae (31.94%) |
| | | Rotavirus A (11.95%) |
| GISAID | 13,859 (3.50%) | Orthopneumovirus hominis (59.20%) |
| | | Orthoflavivirus denguei (20.54%) |
| | | Chikungunya virus (9.02%) |
| LANL-HIV DB | 9,753 (2.46%) | Human immunodeficiency virus 1 (99.89%) |
| | | Simian immunodeficiency virus (0.11%) |
|  |  | ---- |
| HBVdb | 2,728 (0.69%) | Hepatitis B virus (100%) |
| | | --- |
| | | --- |

**Supplementary Note 5**

Description of all datasets used in vGUE Benchmark. The task datasets assess frozen representations, but the accession-based exclusions described below do not, on their own, establish homology-independent train/test separation. Near-ceiling tasks may also have limited ability to differentiate models.

- **Virus vs non-virus classification:** 15000 viral sequences sampled from the test set with the same source-label proportions as the test set (all 295 retained taxonomic labels are represented in this 15,000-sequence sample); for bacteria we sampled 9600 5kb contigs from representative bacterial complete genomes from NCBI RefSeq.

- **Virus vs non-virus reads classification:** Real, non-simulated Illumina paired-end reads sourced from NCBI BioProject PRJNA389927 (deposited on SRA/ENA), originally generated by Hannigan et al. (2018) [42] in their study on the colorectal cancer virome. Inspired by the fraction-based viral-versus-microbial comparison framework of Wu et al. (2024) [S1], two experimental classes were defined for this study by sampling 1,000 reads from specific runs within the project: the viral class was derived from the viral-enriched fraction (SRR5665153), while the putative non-viral/microbial comparator class was derived from the microbial-enriched fraction (SRR5665119). Enrichment of the source fractions does not independently verify a read-level viral/non-viral label for every sequence.

- **DNA vs RNA classification:** 5,000 DNA and 5,000 RNA complete genomes randomly sampled from corpus test set.

- **Host Prediction:** ~2,000 complete viral genomes per class, sampled from the test set across diverse families and organized into four main categories based on host type: vertebrates, invertebrates, plants, and bacteria (phages). To ensure balanced taxonomic coverage, representative families and taxa were selected for each host group. The vertebrate class includes sequences from *Adenoviridae*, *Coronaviridae*, *Orthomyxoviridae*, *Papillomaviridae*, *Paramyxoviridae*, *Parvoviridae*, and *Picornaviridae*. The invertebrate class covers *Spinareoviridae*, *Parvoviridae*, *Iflaviridae*, and *Phasmaviridae*. The plant class incorporates a broad selection including *Bromoviridae*, *Betaflexiviridae*, *Potyviridae*, *Nanoviridae*, *Geminiviridae*, *Spinareoviridae*, *Secoviridae*, *Tospoviridae*, *Rhabdoviridae*, *Fimoviridae*, *Virgaviridae*, *Alphaflexiviridae*, and *Closteroviridae*. Finally, the bacterial phage class is represented by *Caudoviricetes* and *Microviridae*.

- **HIV-1 vs HIV-2 classification: 7**,248 HIV-1 and 94 HIV-2 complete genomes retrieved from the LANL HIV database, **c**orresponding to complete sequences absent from the Vir2vec training corpus according to accession-number cross-referencing. This check does not, by itself, exclude closely homologous sequences.

- **SARS-CoV-2 Lineage Identification:** ~1,000 complete SARS-CoV-2 genomes sampled from GISAID, cross-referenced by accession number to exclude matching training accessions, without establishing a sequence-identity-based holdout. The collection spans seven former Variants of Concern (VOCs) and Variants of Interest (VOIs): Alpha GRY (B.1.1.7+Q.*, first detected in the UK), Beta GH/501Y.V2 (B.1.351+B.1.351.2+B.1.351.3, South Africa), Gamma GR/501Y.V3 (P.1+P.1.*, Brazil/Japan), Delta GK (B.1.617.2+AY.*, India), Omicron GRA (B.1.1.529+BA.*, Botswana/Hong Kong/South Africa), Lambda GR/452Q.V1 (C.37+C.37.1, Peru), and Mu GH (B.1.621+B.1.621.1, Colombia).

- **HIV-1 Tropism Prediction:** 764 brain-derived and 1,325 plasma-derived HIV-1 subtype B env coding sequences (CDS) retrieved from the LANL HIV database. Filtering parameters restricted sequences to subtype B and the env CDS region, strictly excluded ambiguous bases (0% non-ACGT characters), and enforced a single-sequence-per-patient threshold to eliminate patient-level duplication.

**Supplementary Note 6**

Max pooling is selected over mean pooling in **Vir2vec** to prevent the dilution of localized functional motifs and single-nucleotide variants (SNVs) across expanding genomic sequence contexts. In viral genomics, biological functionality and drug-resistance phenotypes are frequently driven by point mutations. Mean pooling averages feature representations across all sequence positions, causing localized mutational signatures to be suppressed by uninformative flanking background nucleotides as sequence window length increases.

To empirically evaluate pooling dynamics, a dilutional benchmark was conducted using the HIV-1 HXB2 reference genome (NC_001802.1). Three key drug-resistance mutation loci were selected:

- **PR_V82A** (Protease, position 2475, T → C)
- **RT_K103N** (Reverse Transcriptase, position 2746, A → T)
- **RT_M184V** (Reverse Transcriptase, position 3098, A → G)

For each locus, sequence representations were extracted across symmetric window lengths $L \in \{256, 512, 1024, 2048, 4096\}$ bp under both max-pooling and mean-pooling architectures. Two primary metrics were measured:

1. **Localized Motif Signal Retention:** Cosine similarity between the sequence embedding of window L and an isolated 41-bp local motif centered on the target locus ($p \pm 20$ bp).
2. **Single-Nucleotide Mutation Sensitivity (ΔSNV):** The $L_2$ norm distance between wild-type and mutant sequence embeddings ($||E\_WT - E\_MUT||_2$).

**Figure S1** shows the dilutional benchmark dynamics across sequence window lengths L, showing (Panel A) localized motif cosine similarity retention and (Panel B) single-nucleotide mutation sensitivity (ΔSNV).

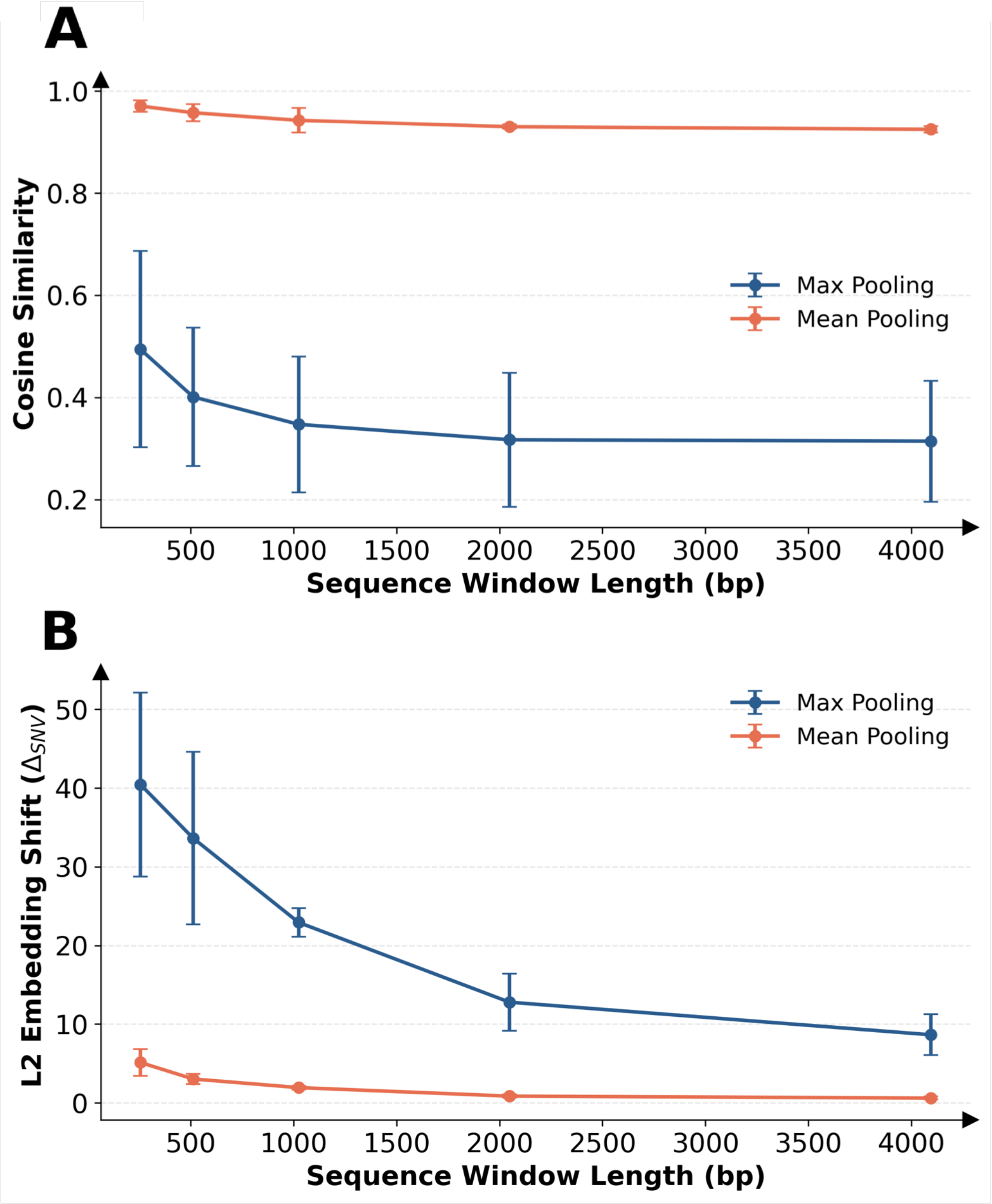


**Fig. S1. Dilutional benchmark dynamics across sequence window lengths L, showing (Panel A) localized motif cosine similarity retention and (Panel B) single-nucleotide mutation sensitivity (ΔSNV).** Mean pooling exhibits higher cosine similarity (≥ 0.925 at 4096 bp), which may reflect broad sequence-compositional similarity rather than specific retention of the local mutation-associated motif. Max pooling yields lower cosine similarity (0.314 at 4096 bp); this metric alone cannot establish superior retention of local motifs. As sequence window length increases to 4096 bp, mean pooling suffers severe signal attenuation, dropping SNV sensitivity to an $L_2$ shift of ΔSNV = 0.61 ± 0.19. Max pooling preserves high-activation feature maps, maintaining an $L_2$ shift of ΔSNV = 8.65 ± 2.60 at 4096 bp—corresponding to a >14-fold larger raw embedding-distance response than mean pooling. Because unnormalised $L_2$ distances can depend on embedding scale, this result does not by itself establish superior downstream mutation detection or sensitivity across full-length viral genomes.

**Table S6. Downstream classification performance on Virus vs Non Virus task from vGUE benchmark across evaluated genomic language models.** Metrics are reported as mean ± standard deviation across 5-fold cross-validation. Evaluated downstream classifiers include One-Class SVM (OCSVM), Isolation Forest (IF) and an Ensemble model combining both. **Metrics shown in the table are Accuracy (Acc), Balanced Accuracy (B.Acc), Macro Precision (Prec), Macro Recall (Rec), Matthews Correlation Coefficient (MCC), and Area Under the Receiver Operating Characteristic Curve (AUROC). Best performance per metric is highlighted in bold.**

| Task | Model | Classifier | Acc | B.Acc | Prec | Rec | MCC | AUROC |
|---|---|---|---|---|---|---|---|---|
| Virus vs Non Virus | Mistral-DNA-v1-138M-hg38 | Ensemble | 0.57 ± 0.13 | 0.50 ± 0.07 | 0.55 ± 0.15 | 0.50 ± 0.07 | 0.03 ± 0.20 | 0.64 ± 0.17 |
| | | IF | 0.58 ± 0.07 | 0.47 ± 0.06 | 0.30 ± 0.02 | 0.47 ± 0.06 | -0.08 ± 0.15 | 0.51 ± 0.23 |
| | | OCSVM | 0.66 ± 0.06 | 0.62 ± 0.05 | 0.65 ± 0.07 | 0.62 ± 0.05 | 0.27 ± 0.12 | 0.66 ± 0.10 |
| | Mistral-DNA-v1-17M-hg38 | Ensemble | 0.57 ± 0.08 | 0.47 ± 0.06 | 0.30 ± 0.02 | 0.47 ± 0.06 | -0.08 ± 0.16 | 0.35 ± 0.17 |
| | | IF | 0.57 ± 0.08 | 0.47 ± 0.06 | 0.30 ± 0.02 | 0.47 ± 0.06 | -0.08 ± 0.16 | 0.34 ± 0.16 |
| | | OCSVM | 0.72 ± 0.10 | 0.71 ± 0.07 | 0.74 ± 0.12 | 0.71 ± 0.07 | 0.45 ± 0.19 | 0.75 ± 0.07 |
| | Mistral-DNA-v1-422M-hg38 | Ensemble | 0.61 ± 0.00 | 0.50 ± 0.00 | 0.43 ± 0.21 | 0.50 ± 0.00 | 0.01 ± 0.05 | 0.40 ± 0.12 |
| | | IF | 0.61 ± 0.01 | 0.50 ± 0.01 | 0.31 ± 0.02 | 0.50 ± 0.01 | -0.03 ± 0.04 | 0.35 ± 0.08 |
| | | OCSVM | 0.73 ± 0.09 | 0.70 ± 0.07 | 0.74 ± 0.12 | 0.70 ± 0.07 | 0.43 ± 0.19 | 0.74 ± 0.08 |
| | ModernBert-DNA-v1-37M-virus | Ensemble | 0.72 ± 0.03 | 0.67 ± 0.03 | 0.76 ± 0.07 | 0.67 ± 0.03 | 0.42 ± 0.07 | 0.72 ± 0.16 |
| | | IF | 0.57 ± 0.08 | 0.47 ± 0.06 | 0.30 ± 0.02 | 0.47 ± 0.06 | -0.08 ± 0.16 | 0.50 ± 0.28 |
| | | OCSVM | 0.73 ± 0.08 | 0.70 ± 0.06 | 0.74 ± 0.11 | 0.70 ± 0.06 | 0.44 ± 0.17 | 0.73 ± 0.08 |
| | Vir2vec-138M | Ensemble | 0.92 ± 0.08 | 0.93 ± 0.07 | 0.92 ± 0.07 | 0.93 ± 0.07 | 0.86 ± 0.14 | 0.95 ± 0.05 |
| | | IF | 0.91 ± 0.09 | 0.92 ± 0.07 | 0.92 ± 0.08 | 0.92 ± 0.07 | 0.84 ± 0.15 | 0.95 ± 0.06 |
| | | OCSVM | 0.60 ± 0.11 | 0.53 ± 0.09 | 0.61 ± 0.16 | 0.53 ± 0.09 | 0.12 ± 0.21 | 0.59 ± 0.28 |
| | Vir2vec-17M | Ensemble | 0.82 ± 0.10 | 0.84 ± 0.08 | 0.83 ± 0.08 | 0.84 ± 0.08 | 0.67 ± 0.16 | 0.91 ± 0.08 |
| | | IF | 0.82 ± 0.10 | 0.83 ± 0.08 | 0.83 ± 0.08 | 0.83 ± 0.08 | 0.67 ± 0.16 | 0.91 ± 0.08 |
| | | OCSVM | 0.70 ± 0.14 | 0.70 ± 0.11 | 0.73 ± 0.14 | 0.70 ± 0.11 | 0.43 ± 0.25 | 0.79 ± 0.14 |
| | Vir2vec-422M | Ensemble | 0.93 ± 0.05 | **0.94 ± 0.04** | **0.93 ± 0.05** | **0.94 ± 0.04** | **0.87 ± 0.09** | **0.97 ± 0.04** |
| | | IF | 0.93 ± 0.06 | 0.93 ± 0.05 | **0.93 ± 0.06** | 0.93 ± 0.05 | 0.86 ± 0.11 | 0.96 ± 0.04 |
| | | OCSVM | 0.63 ± 0.16 | 0.60 ± 0.15 | 0.64 ± 0.19 | 0.60 ± 0.15 | 0.24 ± 0.33 | 0.70 ± 0.25 |
| | evo-1-131k-base | Ensemble | 0.85 ± 0.02 | 0.84 ± 0.03 | 0.86 ± 0.04 | 0.84 ± 0.03 | 0.69 ± 0.06 | 0.84 ± 0.05 |
| | | IF | 0.61 ± 0.00 | 0.50 ± 0.00 | 0.30 ± 0.00 | 0.50 ± 0.00 | -0.02 ± 0.02 | 0.24 ± 0.04 |
| | | OCSVM | 0.85 ± 0.04 | 0.84 ± 0.04 | 0.85 ± 0.05 | 0.84 ± 0.04 | 0.69 ± 0.08 | 0.85 ± 0.05 |
| | evo-1-8k-base | Ensemble | **0.93 ± 0.02** | 0.92 ± 0.02 | **0.93 ± 0.03** | 0.92 ± 0.02 | 0.86 ± 0.05 | 0.94 ± 0.01 |
| | | IF | 0.60 ± 0.00 | 0.50 ± 0.00 | 0.30 ± 0.00 | 0.50 ± 0.00 | -0.05 ± 0.03 | 0.19 ± 0.02 |
| | | OCSVM | 0.92 ± 0.05 | 0.92 ± 0.04 | 0.92 ± 0.05 | 0.92 ± 0.04 | 0.84 ± 0.09 | 0.95 ± 0.01 |

**Table S7. Downstream classification performance on Virus vs Non Virus (Reads) task from vGUE benchmark across evaluated genomic language models.** Metrics are reported as mean ± standard deviation across 5-fold cross-validation. Evaluated downstream classifiers include Logistic Regression (LogReg), Random Forest (RF), and XGBoost (XGB). Metrics shown in the table are Accuracy (Acc), Balanced Accuracy (B.Acc), Macro Precision (Prec), Macro Recall (Rec), Matthews Correlation Coefficient (MCC), and Area Under the Receiver Operating Characteristic Curve (AUROC). Best performance per metric is highlighted in bold.

| Task | Model | Classifier | Acc | B.Acc | Prec | Rec | MCC | AUROC |
|---|---|---|---|---|---|---|---|---|
| Virus vs Non Virus (Reads) | Mistral-DNA-v1-138M-hg38 | LogReg | 0.62 ± 0.02 | 0.62 ± 0.02 | 0.62 ± 0.02 | 0.62 ± 0.02 | 0.25 ± 0.04 | 0.67 ± 0.01 |
| | | RF | - | - | - | - | - | - |
| | | XGB | 0.62 ± 0.03 | 0.62 ± 0.03 | 0.62 ± 0.03 | 0.62 ± 0.03 | 0.23 ± 0.06 | 0.67 ± 0.02 |
| | Mistral-DNA-v1-17M-hg38 | LogReg | 0.70 ± 0.02 | 0.70 ± 0.02 | 0.70 ± 0.02 | 0.70 ± 0.02 | 0.40 ± 0.04 | 0.77 ± 0.03 |
| | | RF | - | - | - | - | - | - |
| | | XGB | **0.71 ± 0.02** | **0.71 ± 0.02** | **0.71 ± 0.02** | **0.71 ± 0.02** | **0.41 ± 0.05** | **0.79 ± 0.02** |
| | Mistral-DNA-v1-422M-hg38 | LogReg | 0.69 ± 0.02 | 0.69 ± 0.02 | 0.69 ± 0.02 | 0.69 ± 0.02 | 0.38 ± 0.04 | 0.77 ± 0.02 |
| | | RF | 0.70 ± 0.03 | 0.70 ± 0.03 | 0.70 ± 0.03 | 0.70 ± 0.03 | 0.41 ± 0.06 | 0.78 ± 0.03 |
| | | XGB | 0.70 ± 0.03 | 0.70 ± 0.03 | 0.70 ± 0.03 | 0.70 ± 0.03 | 0.40 ± 0.05 | 0.78 ± 0.03 |
| | ModernBert-DNA-v1-37M-virus | LogReg | 0.66 ± 0.02 | 0.66 ± 0.02 | 0.66 ± 0.02 | 0.66 ± 0.02 | 0.32 ± 0.05 | 0.72 ± 0.02 |
| | | RF | 0.67 ± 0.02 | 0.67 ± 0.02 | 0.67 ± 0.02 | 0.67 ± 0.02 | 0.34 ± 0.04 | 0.74 ± 0.02 |
| | | XGB | 0.66 ± 0.01 | 0.66 ± 0.01 | 0.66 ± 0.01 | 0.66 ± 0.01 | 0.33 ± 0.03 | 0.72 ± 0.02 |
| | Vir2vec-138M | LogReg | 0.67 ± 0.03 | 0.67 ± 0.03 | 0.67 ± 0.03 | 0.67 ± 0.03 | 0.35 ± 0.06 | 0.73 ± 0.02 |
| | | RF | - | - | - | - | - | - |
| | | XGB | 0.68 ± 0.02 | 0.68 ± 0.02 | 0.68 ± 0.02 | 0.68 ± 0.02 | 0.36 ± 0.03 | 0.74 ± 0.02 |
| | Vir2vec-17M | LogReg | 0.63 ± 0.03 | 0.63 ± 0.03 | 0.63 ± 0.03 | 0.63 ± 0.03 | 0.26 ± 0.06 | 0.67 ± 0.02 |
| | | RF | 0.66 ± 0.01 | 0.66 ± 0.01 | 0.66 ± 0.01 | 0.66 ± 0.01 | 0.31 ± 0.02 | 0.70 ± 0.01 |
| | | XGB | 0.65 ± 0.01 | 0.65 ± 0.01 | 0.65 ± 0.01 | 0.65 ± 0.01 | 0.29 ± 0.03 | 0.69 ± 0.02 |
| | Vir2vec-422M | LogReg | 0.66 ± 0.02 | 0.66 ± 0.02 | 0.66 ± 0.02 | 0.66 ± 0.02 | 0.32 ± 0.03 | 0.72 ± 0.02 |
| | | RF | 0.66 ± 0.02 | 0.66 ± 0.02 | 0.66 ± 0.02 | 0.66 ± 0.02 | 0.32 ± 0.04 | 0.71 ± 0.02 |
| | | XGB | 0.66 ± 0.01 | 0.66 ± 0.01 | 0.66 ± 0.01 | 0.66 ± 0.01 | 0.32 ± 0.02 | 0.72 ± 0.01 |
| | evo-1-131k-base | LogReg | 0.67 ± 0.03 | 0.67 ± 0.03 | 0.67 ± 0.03 | 0.67 ± 0.03 | 0.34 ± 0.05 | 0.73 ± 0.03 |
| | | RF | 0.68 ± 0.02 | 0.68 ± 0.02 | 0.68 ± 0.02 | 0.68 ± 0.02 | 0.35 ± 0.04 | 0.77 ± 0.02 |
| | | XGB | 0.68 ± 0.02 | 0.68 ± 0.02 | 0.68 ± 0.02 | 0.68 ± 0.02 | 0.37 ± 0.04 | 0.77 ± 0.03 |
| | evo-1-8k-base | LogReg | 0.62 ± 0.03 | 0.62 ± 0.03 | 0.62 ± 0.03 | 0.62 ± 0.03 | 0.24 ± 0.05 | 0.67 ± 0.02 |
| | | RF | 0.70 ± 0.02 | 0.70 ± 0.02 | 0.70 ± 0.02 | 0.70 ± 0.02 | 0.39 ± 0.04 | 0.78 ± 0.02 |
| | | XGB | 0.69 ± 0.02 | 0.69 ± 0.02 | 0.69 ± 0.02 | 0.69 ± 0.02 | 0.38 ± 0.03 | 0.76 ± 0.02 |

**Table S8. Downstream classification performance on DNA vs RNA task from vGUE benchmark across evaluated genomic language models.** Metrics are reported as mean ± standard deviation across 5-fold cross-validation. Evaluated downstream classifiers include Logistic Regression (LogReg), Random Forest (RF), and XGBoost (XGB). Metrics shown in the table are Accuracy (Acc), Balanced Accuracy (B.Acc), Macro Precision (Prec), Macro Recall (Rec), Matthews Correlation Coefficient (MCC), and Area Under the Receiver Operating Characteristic Curve (AUROC). Best performance per metric is highlighted in bold.

| Task | Model | Classifier | Acc | B.Acc | Prec | Rec | MCC | AUROC |
|---|---|---|---|---|---|---|---|---|
| DNA vs RNA | Mistral-DNA-v1-138M-hg38 | LogReg | 0.83 ± 0.00 | 0.83 ± 0.00 | 0.83 ± 0.00 | 0.83 ± 0.00 | 0.66 ± 0.01 | 0.90 ± 0.01 |
| | | RF | 0.89 ± 0.01 | 0.89 ± 0.01 | 0.90 ± 0.01 | 0.89 ± 0.01 | 0.79 ± 0.01 | 0.94 ± 0.01 |
| | | XGB | 0.90 ± 0.01 | 0.90 ± 0.01 | 0.90 ± 0.01 | 0.90 ± 0.01 | 0.80 ± 0.02 | 0.95 ± 0.00 |
| | Mistral-DNA-v1-17M-hg38 | LogReg | 0.85 ± 0.01 | 0.85 ± 0.01 | 0.85 ± 0.01 | 0.85 ± 0.01 | 0.70 ± 0.02 | 0.92 ± 0.01 |
| | | RF | 0.93 ± 0.00 | 0.93 ± 0.00 | 0.93 ± 0.00 | 0.93 ± 0.00 | 0.86 ± 0.01 | 0.98 ± 0.00 |
| | | XGB | 0.93 ± 0.00 | 0.93 ± 0.00 | 0.93 ± 0.00 | 0.93 ± 0.00 | 0.85 ± 0.01 | 0.98 ± 0.00 |
| | Mistral-DNA-v1-422M-hg38 | LogReg | 0.91 ± 0.01 | 0.91 ± 0.01 | 0.92 ± 0.01 | 0.91 ± 0.01 | 0.83 ± 0.01 | 0.97 ± 0.01 |
| | | RF | 0.90 ± 0.01 | 0.91 ± 0.01 | 0.91 ± 0.01 | 0.91 ± 0.01 | 0.82 ± 0.01 | 0.96 ± 0.00 |
| | | XGB | 0.91 ± 0.01 | 0.91 ± 0.01 | 0.92 ± 0.01 | 0.91 ± 0.01 | 0.83 ± 0.01 | 0.97 ± 0.00 |
| | ModernBert-DNA-v1-37M-virus | LogReg | 0.88 ± 0.01 | 0.88 ± 0.01 | 0.88 ± 0.01 | 0.88 ± 0.01 | 0.77 ± 0.01 | 0.94 ± 0.01 |
| | | RF | 0.87 ± 0.01 | 0.88 ± 0.01 | 0.89 ± 0.01 | 0.88 ± 0.01 | 0.77 ± 0.02 | 0.93 ± 0.01 |
| | | XGB | 0.89 ± 0.01 | 0.89 ± 0.01 | 0.90 ± 0.01 | 0.89 ± 0.01 | 0.78 ± 0.02 | 0.95 ± 0.01 |
| | Vir2vec-138M | LogReg | 0.91 ± 0.01 | 0.91 ± 0.01 | 0.91 ± 0.01 | 0.91 ± 0.01 | 0.82 ± 0.02 | 0.96 ± 0.01 |
| | | RF | 0.94 ± 0.01 | 0.94 ± 0.01 | 0.94 ± 0.01 | 0.94 ± 0.01 | 0.88 ± 0.01 | 0.98 ± 0.00 |
| | | XGB | 0.94 ± 0.01 | 0.94 ± 0.01 | 0.94 ± 0.01 | 0.94 ± 0.01 | 0.88 ± 0.01 | 0.99 ± 0.00 |
| | Vir2vec-17M | LogReg | 0.88 ± 0.01 | 0.88 ± 0.01 | 0.88 ± 0.01 | 0.88 ± 0.01 | 0.76 ± 0.02 | 0.94 ± 0.01 |
| | | RF | 0.94 ± 0.00 | 0.94 ± 0.00 | 0.94 ± 0.00 | 0.94 ± 0.00 | 0.88 ± 0.01 | **0.99 ± 0.00** |
| | | XGB | **0.95 ± 0.01** | **0.95 ± 0.01** | **0.95 ± 0.01** | **0.95 ± 0.01** | **0.89 ± 0.01** | **0.99 ± 0.00** |
| | Vir2vec-422M | LogReg | 0.90 ± 0.01 | 0.90 ± 0.01 | 0.90 ± 0.01 | 0.90 ± 0.01 | 0.80 ± 0.02 | 0.96 ± 0.01 |
| | | RF | 0.94 ± 0.01 | 0.94 ± 0.01 | 0.94 ± 0.01 | 0.94 ± 0.01 | 0.88 ± 0.01 | 0.98 ± 0.00 |
| | | XGB | 0.94 ± 0.01 | 0.94 ± 0.01 | 0.94 ± 0.01 | 0.94 ± 0.01 | 0.88 ± 0.01 | **0.99 ± 0.00** |
| | evo-1-131k-base | LogReg | 0.93 ± 0.01 | 0.93 ± 0.01 | 0.93 ± 0.01 | 0.93 ± 0.01 | 0.86 ± 0.01 | 0.97 ± 0.00 |
| | | RF | 0.94 ± 0.01 | 0.94 ± 0.01 | 0.94 ± 0.01 | 0.94 ± 0.01 | **0.89 ± 0.01** | 0.98 ± 0.00 |
| | | XGB | 0.94 ± 0.00 | 0.94 ± 0.00 | 0.94 ± 0.00 | 0.94 ± 0.00 | 0.88 ± 0.01 | 0.98 ± 0.00 |
| | evo-1-8k-base | LogReg | 0.91 ± 0.01 | 0.91 ± 0.01 | 0.91 ± 0.01 | 0.91 ± 0.01 | 0.82 ± 0.01 | 0.96 ± 0.01 |
| | | RF | 0.93 ± 0.01 | 0.93 ± 0.01 | 0.93 ± 0.01 | 0.93 ± 0.01 | 0.86 ± 0.01 | 0.97 ± 0.00 |
| | | XGB | 0.92 ± 0.01 | 0.92 ± 0.01 | 0.92 ± 0.01 | 0.92 ± 0.01 | 0.84 ± 0.02 | 0.97 ± 0.01 |

**Table S9. Downstream classification performance on HIV-1 vs HIV-2 task from vGUE benchmark across evaluated genomic language models.** Metrics are reported as mean ± standard deviation across 5-fold cross-validation. Evaluated downstream classifiers include Logistic Regression (LogReg), Random Forest (RF), and XGBoost (XGB). Metrics shown in the table are Accuracy (Acc), Balanced Accuracy (B.Acc), Macro Precision (Prec), Macro Recall (Rec), Matthews Correlation Coefficient (MCC), and Area Under the Receiver Operating Characteristic Curve (AUROC). Best performance per metric is highlighted in bold.

| Task | Model | Classifier | Acc | B.Acc | Prec | Rec | MCC | AUROC |
|---|---|---|---|---|---|---|---|---|
| HIV-1 vs HIV-2 | Mistral-DNA-v1-138M-hg38 | LogReg | **1.00 ± 0.00** | **1.00 ± 0.00** | 0.89 ± 0.02 | **1.00 ± 0.00** | 0.89 ± 0.03 | **1.00 ± 0.00** |
| | | RF | **1.00 ± 0.00** | 0.81 ± 0.02 | **1.00 ± 0.00** | 0.81 ± 0.02 | 0.79 ± 0.03 | 0.99 ± 0.01 |
| | | XGB | **1.00 ± 0.00** | 0.87 ± 0.03 | 0.99 ± 0.01 | 0.87 ± 0.03 | 0.86 ± 0.02 | 0.99 ± 0.01 |
| | Mistral-DNA-v1-17M-hg38 | LogReg | 0.99 ± 0.00 | 0.82 ± 0.07 | 0.76 ± 0.03 | 0.82 ± 0.07 | 0.57 ± 0.08 | 0.94 ± 0.02 |
| | | RF | 0.99 ± 0.00 | 0.59 ± 0.04 | 0.90 ± 0.10 | 0.59 ± 0.04 | 0.36 ± 0.07 | 0.80 ± 0.04 |
| | | XGB | 0.99 ± 0.00 | 0.52 ± 0.01 | 0.79 ± 0.27 | 0.52 ± 0.01 | 0.14 ± 0.12 | 0.92 ± 0.05 |
| | Mistral-DNA-v1-422M-hg38 | LogReg | **1.00 ± 0.00** | **1.00 ± 0.00** | 0.99 ± 0.01 | **1.00 ± 0.00** | 0.99 ± 0.01 | **1.00 ± 0.00** |
| | | RF | **1.00 ± 0.00** | 0.92 ± 0.07 | 0.99 ± 0.01 | 0.92 ± 0.07 | 0.91 ± 0.07 | 0.99 ± 0.01 |
| | | XGB | **1.00 ± 0.00** | 0.95 ± 0.02 | 0.99 ± 0.02 | 0.95 ± 0.02 | 0.93 ± 0.03 | **1.00 ± 0.00** |
| | ModernBert-DNA-v1-37M-virus | LogReg | **1.00 ± 0.00** | **1.00 ± 0.00** | 0.97 ± 0.02 | **1.00 ± 0.00** | 0.97 ± 0.02 | **1.00 ± 0.00** |
| | | RF | **1.00 ± 0.00** | 0.91 ± 0.02 | 0.99 ± 0.01 | 0.91 ± 0.02 | 0.90 ± 0.03 | **1.00 ± 0.00** |
| | | XGB | **1.00 ± 0.00** | 0.93 ± 0.03 | 0.99 ± 0.02 | 0.93 ± 0.03 | 0.91 ± 0.05 | **1.00 ± 0.00** |
| | Vir2vec-138M | LogReg | **1.00 ± 0.00** | **1.00 ± 0.00** | **1.00 ± 0.00** | 1.00 ± 0.00 | **1.00 ± 0.00** | **1.00 ± 0.00** |
| | | RF | **1.00 ± 0.00** | 0.99 ± 0.01 | 0.97 ± 0.04 | 0.99 ± 0.01 | 0.96 ± 0.03 | **1.00 ± 0.00** |
| | | XGB | **1.00 ± 0.00** | 0.99 ± 0.01 | 0.99 ± 0.01 | 0.99 ± 0.01 | 0.98 ± 0.01 | **1.00 ± 0.00** |
| | Vir2vec-17M | LogReg | **1.00 ± 0.00** | 0.99 ± 0.01 | 0.98 ± 0.02 | 0.99 ± 0.01 | 0.97 ± 0.02 | **1.00 ± 0.00** |
| | | RF | **1.00 ± 0.00** | **1.00 ± 0.00** | 0.98 ± 0.02 | **1.00 ± 0.00** | 0.98 ± 0.02 | **1.00 ± 0.00** |
| | | XGB | **1.00 ± 0.00** | **1.00 ± 0.00** | 0.94 ± 0.03 | **1.00 ± 0.00** | 0.93 ± 0.04 | **1.00 ± 0.00** |
| | Vir2vec-422M | LogReg | **1.00 ± 0.00** | **1.00 ± 0.00** | 0.99 ± 0.02 | **1.00 ± 0.00** | 0.99 ± 0.02 | **1.00 ± 0.00** |
| | | RF | **1.00 ± 0.00** | 0.99 ± 0.01 | 0.99 ± 0.01 | 0.99 ± 0.01 | 0.99 ± 0.01 | **1.00 ± 0.00** |
| | | XGB | **1.00 ± 0.00** | **1.00 ± 0.00** | 0.99 ± 0.01 | 1.00 ± 0.00 | 0.99 ± 0.01 | **1.00 ± 0.00** |
| | evo-1-131k-base | LogReg | 0.99 ± 0.00 | 0.98 ± 0.01 | 0.79 ± 0.05 | 0.98 ± 0.01 | 0.75 ± 0.06 | **1.00 ± 0.00** |
| | | RF | 0.99 ± 0.00 | 0.87 ± 0.06 | 0.90 ± 0.07 | 0.87 ± 0.06 | 0.76 ± 0.09 | 0.99 ± 0.02 |
| | | XGB | 0.99 ± 0.00 | 0.92 ± 0.02 | 0.87 ± 0.04 | 0.92 ± 0.02 | 0.79 ± 0.03 | **1.00 ± 0.01** |
| | evo-1-8k-base | LogReg | 0.91 ± 0.01 | 0.81 ± 0.08 | 0.55 ± 0.01 | 0.81 ± 0.08 | 0.24 ± 0.05 | 0.91 ± 0.02 |
| | | RF | 0.99 ± 0.00 | 0.70 ± 0.03 | 0.72 ± 0.04 | 0.70 ± 0.03 | 0.41 ± 0.03 | 0.92 ± 0.03 |
| | | XGB | 0.99 ± 0.00 | 0.71 ± 0.07 | 0.72 ± 0.07 | 0.71 ± 0.07 | 0.42 ± 0.12 | 0.91 ± 0.03 |

**Table S10. Downstream classification performance on SARS-CoV-2 Variant Identification task from vGUE benchmark across evaluated genomic language models.** Metrics are reported as mean ± standard deviation across 5-fold cross-validation. Evaluated downstream classifiers include Logistic Regression (LogReg), Random Forest (RF), and XGBoost (XGB). Metrics shown in the table are Accuracy (Acc), Balanced Accuracy (B.Acc), Macro Precision (Prec), Macro Recall (Rec), Matthews Correlation Coefficient (MCC), and Area Under the Receiver Operating Characteristic Curve (AUROC). Best performance per metric is highlighted in bold.

| Task | Model | Classifier | Acc | B.Acc | Prec | Rec | MCC | AUROC |
|---|---|---|---|---|---|---|---|---|
| SARS-CoV-2 Variant Identification | Mistral-DNA-v1-138M-hg38 | LogReg | **0.99 ± 0.00** | **0.99 ± 0.00** | **0.99 ± 0.00** | **0.99 ± 0.00** | **0.99 ± 0.00** | **1.00 ± 0.00** |
| | | RF | 0.95 ± 0.01 | 0.95 ± 0.01 | 0.95 ± 0.01 | 0.95 ± 0.01 | 0.95 ± 0.01 | **1.00 ± 0.00** |
| | | XGB | 0.97 ± 0.01 | 0.97 ± 0.01 | 0.97 ± 0.01 | 0.97 ± 0.01 | 0.97 ± 0.01 | **1.00 ± 0.00** |
| | Mistral-DNA-v1-17M-hg38 | LogReg | 0.92 ± 0.01 | 0.92 ± 0.01 | 0.92 ± 0.01 | 0.92 ± 0.01 | 0.91 ± 0.01 | 0.98 ± 0.00 |
| | | RF | 0.91 ± 0.00 | 0.91 ± 0.00 | 0.91 ± 0.00 | 0.91 ± 0.00 | 0.89 ± 0.00 | 0.99 ± 0.00 |
| | | XGB | 0.93 ± 0.01 | 0.93 ± 0.01 | 0.93 ± 0.01 | 0.93 ± 0.01 | 0.92 ± 0.01 | **1.00 ± 0.00** |
| | Mistral-DNA-v1-422M-hg38 | LogReg | 0.97 ± 0.01 | 0.97 ± 0.01 | 0.97 ± 0.01 | 0.97 ± 0.01 | 0.96 ± 0.01 | **1.00 ± 0.00** |
| | | RF | 0.93 ± 0.01 | 0.93 ± 0.01 | 0.93 ± 0.01 | 0.93 ± 0.01 | 0.92 ± 0.01 | 0.99 ± 0.00 |
| | | XGB | 0.95 ± 0.01 | 0.95 ± 0.01 | 0.95 ± 0.01 | 0.95 ± 0.01 | 0.94 ± 0.01 | **1.00 ± 0.00** |
| | ModernBert-DNA-v1-37M-virus | LogReg | **0.99 ± 0.00** | **0.99 ± 0.00** | **0.99 ± 0.00** | **0.99 ± 0.00** | 0.98 ± 0.00 | **1.00 ± 0.00** |
| | | RF | 0.95 ± 0.01 | 0.95 ± 0.01 | 0.95 ± 0.01 | 0.95 ± 0.01 | 0.94 ± 0.01 | **1.00 ± 0.00** |
| | | XGB | 0.97 ± 0.00 | 0.97 ± 0.00 | 0.97 ± 0.00 | 0.97 ± 0.00 | 0.97 ± 0.00 | **1.00 ± 0.00** |
| | Vir2vec-138M | LogReg | 0.98 ± 0.00 | 0.98 ± 0.00 | 0.98 ± 0.00 | 0.98 ± 0.00 | 0.98 ± 0.00 | **1.00 ± 0.00** |
| | | RF | 0.94 ± 0.01 | 0.94 ± 0.01 | 0.95 ± 0.01 | 0.94 ± 0.01 | 0.94 ± 0.01 | 0.99 ± 0.00 |
| | | XGB | 0.96 ± 0.00 | 0.96 ± 0.00 | 0.96 ± 0.00 | 0.96 ± 0.00 | 0.95 ± 0.00 | **1.00 ± 0.00** |
| | Vir2vec-17M | LogReg | 0.98 ± 0.00 | 0.98 ± 0.00 | 0.98 ± 0.00 | 0.98 ± 0.00 | 0.98 ± 0.00 | **1.00 ± 0.00** |
| | | RF | 0.97 ± 0.01 | 0.97 ± 0.01 | 0.97 ± 0.01 | 0.97 ± 0.01 | 0.96 ± 0.01 | **1.00 ± 0.00** |
| | | XGB | 0.97 ± 0.00 | 0.97 ± 0.00 | 0.98 ± 0.00 | 0.97 ± 0.00 | 0.97 ± 0.01 | **1.00 ± 0.00** |
| | Vir2vec-422M | LogReg | 0.98 ± 0.00 | 0.98 ± 0.00 | 0.98 ± 0.00 | 0.98 ± 0.00 | 0.98 ± 0.00 | **1.00 ± 0.00** |
| | | RF | 0.96 ± 0.01 | 0.96 ± 0.01 | 0.96 ± 0.01 | 0.96 ± 0.01 | 0.95 ± 0.01 | **1.00 ± 0.00** |
| | | XGB | 0.97 ± 0.01 | 0.97 ± 0.01 | 0.97 ± 0.00 | 0.97 ± 0.01 | 0.96 ± 0.01 | **1.00 ± 0.00** |
| | evo-1-131k-base | LogReg | 0.85 ± 0.01 | 0.85 ± 0.01 | 0.85 ± 0.01 | 0.85 ± 0.01 | 0.83 ± 0.02 | 0.97 ± 0.00 |
| | | RF | 0.81 ± 0.02 | 0.81 ± 0.02 | 0.81 ± 0.02 | 0.81 ± 0.02 | 0.78 ± 0.03 | 0.97 ± 0.00 |
| | | XGB | 0.84 ± 0.02 | 0.84 ± 0.02 | 0.84 ± 0.02 | 0.84 ± 0.02 | 0.81 ± 0.02 | 0.98 ± 0.00 |
| | evo-1-8k-base | LogReg | 0.71 ± 0.01 | 0.71 ± 0.01 | 0.70 ± 0.01 | 0.71 ± 0.01 | 0.66 ± 0.01 | 0.93 ± 0.01 |
| | | RF | 0.76 ± 0.01 | 0.76 ± 0.01 | 0.76 ± 0.01 | 0.76 ± 0.01 | 0.72 ± 0.01 | 0.95 ± 0.00 |
| | | XGB | 0.75 ± 0.01 | 0.75 ± 0.01 | 0.75 ± 0.01 | 0.75 ± 0.01 | 0.71 ± 0.01 | 0.96 ± 0.00 |

**Table S11. Downstream classification performance on Host Prediction task from vGUE benchmark across evaluated genomic language models.** Metrics are reported as mean ± standard deviation across 5-fold cross-validation. Evaluated downstream classifiers include Logistic Regression (LogReg), Random Forest (RF), and XGBoost (XGB). Metrics shown in the table are Accuracy (Acc), Balanced Accuracy (B.Acc), Macro Precision (Prec), Macro Recall (Rec), Matthews Correlation Coefficient (MCC), and Area Under the Receiver Operating Characteristic Curve (AUROC). Best performance per metric is highlighted in bold.

| Task | Model | Classifier | Acc | B.Acc | Prec | Rec | MCC | AUROC |
|---|---|---|---|---|---|---|---|---|
| Host Prediction | Mistral-DNA-v1-138M-hg38 | LogReg | 0.80 ± 0.01 | 0.80 ± 0.01 | 0.80 ± 0.01 | 0.80 ± 0.01 | 0.73 ± 0.02 | 0.95 ± 0.00 |
| | | RF | 0.78 ± 0.01 | 0.78 ± 0.01 | 0.78 ± 0.01 | 0.78 ± 0.01 | 0.71 ± 0.02 | 0.93 ± 0.00 |
| | | XGB | 0.81 ± 0.01 | 0.81 ± 0.01 | 0.82 ± 0.01 | 0.81 ± 0.01 | 0.75 ± 0.02 | 0.95 ± 0.00 |
| | Mistral-DNA-v1-17M-hg38 | LogReg | 0.83 ± 0.01 | 0.83 ± 0.01 | 0.83 ± 0.01 | 0.83 ± 0.01 | 0.78 ± 0.01 | 0.96 ± 0.00 |
| | | RF | 0.87 ± 0.00 | 0.87 ± 0.00 | 0.87 ± 0.00 | 0.87 ± 0.00 | 0.83 ± 0.00 | 0.96 ± 0.00 |
| | | XGB | 0.86 ± 0.00 | 0.86 ± 0.00 | 0.87 ± 0.00 | 0.86 ± 0.00 | 0.82 ± 0.00 | 0.97 ± 0.00 |
| | Mistral-DNA-v1-422M-hg38 | LogReg | 0.83 ± 0.01 | 0.83 ± 0.01 | 0.83 ± 0.01 | 0.83 ± 0.01 | 0.77 ± 0.01 | 0.96 ± 0.00 |
| | | RF | 0.84 ± 0.01 | 0.84 ± 0.01 | 0.84 ± 0.01 | 0.84 ± 0.01 | 0.78 ± 0.01 | 0.95 ± 0.00 |
| | | XGB | 0.85 ± 0.01 | 0.85 ± 0.01 | 0.85 ± 0.01 | 0.85 ± 0.01 | 0.80 ± 0.01 | 0.96 ± 0.00 |
| | ModernBert-DNA-v1-37M-virus | LogReg | 0.87 ± 0.00 | 0.87 ± 0.00 | 0.87 ± 0.00 | 0.87 ± 0.00 | 0.82 ± 0.01 | 0.97 ± 0.00 |
| | | RF | 0.85 ± 0.01 | 0.85 ± 0.01 | 0.86 ± 0.01 | 0.85 ± 0.01 | 0.80 ± 0.01 | 0.96 ± 0.00 |
| | | XGB | 0.87 ± 0.01 | 0.87 ± 0.01 | 0.87 ± 0.01 | 0.87 ± 0.01 | 0.82 ± 0.01 | 0.97 ± 0.00 |
| | Vir2vec-138M | LogReg | 0.89 ± 0.01 | 0.89 ± 0.01 | 0.90 ± 0.01 | 0.89 ± 0.01 | 0.86 ± 0.01 | **0.98 ± 0.00** |
| | | RF | 0.90 ± 0.00 | **0.90 ± 0.00** | **0.91 ± 0.00** | **0.90 ± 0.00** | 0.87 ± 0.00 | **0.98 ± 0.00** |
| | | XGB | **0.91 ± 0.00** | **0.90 ± 0.00** | **0.91 ± 0.00** | **0.90 ± 0.00** | **0.88 ± 0.01** | **0.98 ± 0.00** |
| | Vir2vec-17M | LogReg | 0.87 ± 0.01 | 0.87 ± 0.01 | 0.87 ± 0.01 | 0.87 ± 0.01 | 0.83 ± 0.01 | 0.97 ± 0.00 |
| | | RF | 0.87 ± 0.01 | 0.87 ± 0.01 | 0.87 ± 0.01 | 0.87 ± 0.01 | 0.82 ± 0.01 | 0.96 ± 0.00 |
| | | XGB | 0.88 ± 0.01 | 0.88 ± 0.01 | 0.88 ± 0.01 | 0.88 ± 0.01 | 0.84 ± 0.01 | 0.97 ± 0.00 |
| | Vir2vec-422M | LogReg | 0.89 ± 0.01 | 0.89 ± 0.01 | 0.90 ± 0.01 | 0.89 ± 0.01 | 0.86 ± 0.01 | **0.98 ± 0.00** |
| | | RF | 0.90 ± 0.00 | **0.90 ± 0.00** | **0.91 ± 0.00** | **0.90 ± 0.00** | 0.87 ± 0.00 | 0.97 ± 0.00 |
| | | XGB | 0.90 ± 0.01 | **0.90 ± 0.01** | **0.91 ± 0.01** | **0.90 ± 0.01** | 0.87 ± 0.01 | **0.98 ± 0.00** |
| | evo-1-131k-base | LogReg | 0.78 ± 0.01 | 0.78 ± 0.01 | 0.78 ± 0.01 | 0.78 ± 0.01 | 0.70 ± 0.01 | 0.93 ± 0.00 |
| | | RF | 0.87 ± 0.01 | 0.87 ± 0.01 | 0.88 ± 0.01 | 0.87 ± 0.01 | 0.83 ± 0.01 | 0.97 ± 0.00 |
| | | XGB | 0.88 ± 0.01 | 0.88 ± 0.01 | 0.89 ± 0.01 | 0.88 ± 0.01 | 0.85 ± 0.01 | 0.97 ± 0.00 |
| | evo-1-8k-base | LogReg | 0.70 ± 0.01 | 0.70 ± 0.01 | 0.70 ± 0.01 | 0.70 ± 0.01 | 0.60 ± 0.01 | 0.88 ± 0.00 |
| | | RF | 0.84 ± 0.01 | 0.84 ± 0.01 | 0.85 ± 0.01 | 0.84 ± 0.01 | 0.79 ± 0.01 | 0.96 ± 0.00 |
| | | XGB | 0.84 ± 0.01 | 0.84 ± 0.01 | 0.84 ± 0.01 | 0.84 ± 0.01 | 0.78 ± 0.01 | 0.96 ± 0.00 |

**Table S12. Downstream classification performance on HIV-1 Tropism Prediction task from vGUE benchmark across evaluated genomic language models.** Metrics are reported as mean ± standard deviation across 5-fold cross-validation. Evaluated downstream classifiers include Logistic Regression (LogReg), Random Forest (RF), and XGBoost (XGB). Metrics shown in the table are Accuracy (Acc), Balanced Accuracy (B.Acc), Macro Precision (Prec), Macro Recall (Rec), Matthews Correlation Coefficient (MCC), and Area Under the Receiver Operating Characteristic Curve (AUROC). Best performance per metric is highlighted in bold.

| Task | Model | Classifier | Acc | B.Acc | Prec | Rec | MCC | AUROC |
|---|---|---|---|---|---|---|---|---|
| HIV-1 Tropism Prediction | Mistral-DNA-v1-138M-hg38 | LogReg | 0.96 ± 0.01 | 0.96 ± 0.01 | 0.95 ± 0.01 | 0.96 ± 0.01 | 0.91 ± 0.02 | **0.99 ± 0.01** |
| | | RF | 0.93 ± 0.01 | 0.90 ± 0.01 | 0.94 ± 0.01 | 0.90 ± 0.01 | 0.84 ± 0.02 | 0.97 ± 0.01 |
| | | XGB | 0.94 ± 0.01 | 0.92 ± 0.01 | 0.96 ± 0.01 | 0.92 ± 0.01 | 0.88 ± 0.01 | 0.98 ± 0.01 |
| | Mistral-DNA-v1-17M-hg38 | LogReg | 0.96 ± 0.01 | 0.95 ± 0.01 | 0.95 ± 0.01 | 0.95 ± 0.01 | 0.91 ± 0.02 | **0.99 ± 0.00** |
| | | RF | 0.94 ± 0.01 | 0.92 ± 0.01 | 0.95 ± 0.01 | 0.92 ± 0.01 | 0.87 ± 0.02 | 0.98 ± 0.00 |
| | | XGB | 0.95 ± 0.01 | 0.94 ± 0.01 | 0.96 ± 0.01 | 0.94 ± 0.01 | 0.90 ± 0.02 | **0.99 ± 0.01** |
| | Mistral-DNA-v1-422M-hg38 | LogReg | 0.96 ± 0.01 | 0.96 ± 0.01 | 0.96 ± 0.01 | 0.96 ± 0.01 | 0.92 ± 0.02 | **0.99 ± 0.00** |
| | | RF | 0.94 ± 0.01 | 0.92 ± 0.01 | 0.95 ± 0.01 | 0.92 ± 0.01 | 0.87 ± 0.03 | 0.98 ± 0.01 |
| | | XGB | 0.95 ± 0.01 | 0.93 ± 0.01 | 0.96 ± 0.01 | 0.93 ± 0.01 | 0.89 ± 0.02 | **0.99 ± 0.01** |
| | ModernBert-DNA-v1-37M-virus | LogReg | **0.97 ± 0.01** | **0.97 ± 0.01** | **0.97 ± 0.01** | **0.97 ± 0.01** | **0.94 ± 0.02** | **0.99 ± 0.01** |
| | | RF | 0.94 ± 0.01 | 0.92 ± 0.01 | 0.95 ± 0.01 | 0.92 ± 0.01 | 0.87 ± 0.02 | 0.98 ± 0.00 |
| | | XGB | 0.96 ± 0.01 | 0.94 ± 0.01 | **0.97 ± 0.01** | 0.94 ± 0.01 | 0.91 ± 0.02 | **0.99 ± 0.01** |
| | Vir2vec-138M | LogReg | **0.97 ± 0.01** | 0.96 ± 0.01 | **0.97 ± 0.01** | 0.96 ± 0.01 | 0.93 ± 0.02 | **0.99 ± 0.01** |
| | | RF | 0.92 ± 0.02 | 0.90 ± 0.02 | 0.94 ± 0.01 | 0.90 ± 0.02 | 0.83 ± 0.03 | 0.96 ± 0.01 |
| | | XGB | 0.92 ± 0.02 | 0.90 ± 0.02 | 0.95 ± 0.01 | 0.90 ± 0.02 | 0.84 ± 0.03 | 0.98 ± 0.01 |
| | Vir2vec-17M | LogReg | 0.96 ± 0.01 | 0.95 ± 0.01 | 0.96 ± 0.02 | 0.95 ± 0.01 | 0.91 ± 0.03 | **0.99 ± 0.01** |
| | | RF | 0.93 ± 0.01 | 0.91 ± 0.01 | 0.95 ± 0.01 | 0.91 ± 0.01 | 0.85 ± 0.02 | 0.98 ± 0.01 |
| | | XGB | 0.94 ± 0.01 | 0.92 ± 0.01 | 0.96 ± 0.01 | 0.92 ± 0.01 | 0.88 ± 0.02 | **0.99 ± 0.00** |
| | Vir2vec-422M | LogReg | 0.95 ± 0.01 | 0.94 ± 0.01 | 0.95 ± 0.01 | 0.94 ± 0.01 | 0.89 ± 0.02 | **0.99 ± 0.00** |
| | | RF | 0.91 ± 0.01 | 0.88 ± 0.01 | 0.93 ± 0.01 | 0.88 ± 0.01 | 0.80 ± 0.01 | 0.95 ± 0.01 |
| | | XGB | 0.91 ± 0.01 | 0.87 ± 0.01 | 0.93 ± 0.01 | 0.87 ± 0.01 | 0.81 ± 0.02 | 0.96 ± 0.00 |
| | evo-1-131k-base | LogReg | 0.82 ± 0.02 | 0.81 ± 0.02 | 0.80 ± 0.02 | 0.81 ± 0.02 | 0.61 ± 0.04 | 0.90 ± 0.01 |
| | | RF | 0.94 ± 0.02 | 0.92 ± 0.02 | 0.95 ± 0.01 | 0.92 ± 0.02 | 0.86 ± 0.04 | 0.97 ± 0.02 |
| | | XGB | 0.93 ± 0.02 | 0.91 ± 0.02 | 0.94 ± 0.01 | 0.91 ± 0.02 | 0.85 ± 0.03 | 0.97 ± 0.02 |
| | evo-1-8k-base | LogReg | 0.82 ± 0.01 | 0.78 ± 0.01 | 0.83 ± 0.02 | 0.78 ± 0.01 | 0.61 ± 0.02 | 0.82 ± 0.02 |
| | | RF | 0.91 ± 0.01 | 0.89 ± 0.02 | 0.93 ± 0.01 | 0.89 ± 0.02 | 0.82 ± 0.02 | 0.95 ± 0.02 |
| | | XGB | 0.89 ± 0.02 | 0.87 ± 0.02 | 0.89 ± 0.02 | 0.87 ± 0.02 | 0.76 ± 0.03 | 0.94 ± 0.02 |
